# A SARS-CoV-2 Spike Ferritin Nanoparticle Vaccine is Protective and Promotes a Strong Immunological Response in the Cynomolgus Macaque Coronavirus Disease 2019 (COVID-19) Model

**DOI:** 10.1101/2022.03.25.485832

**Authors:** Sara C. Johnston, Keersten M. Ricks, Ines Lakhal-Naouar, Alexandra Jay, Caroline Subra, Jo Lynne Raymond, Hannah A. D. King, Franco Rossi, Tamara L. Clements, David Fetterer, Samantha Tostenson, Camila Macedo Cincotta, Holly R. Hack, Caitlin Kuklis, Sandrine Soman, Jocelyn King, Kristina K. Peachman, Dohoon Kim, Wei-Hung Chen, Rajeshwer S. Sankhala, Elizabeth J. Martinez, Agnes Hajduczki, William C. Chang, Misook Choe, Paul V. Thomas, Caroline E. Peterson, Alexander Anderson, Isabella Swafford, Jeffrey R. Currier, Dominic Paquin-Proulx, Linda L. Jagodzinski, Gary R. Matyas, Mangala Rao, Gregory D. Gromowski, Sheila A. Peel, Lauren White, Jeffrey M. Smith, Jay W. Hooper, Nelson L. Michael, Kayvon Modjarrad, M. Gordon Joyce, Aysegul Nalca, Diane L. Bolton, Margaret LM Pitt

**Author notes:** **Corresponding author:** (SCJ). These authors contributed equally to this work (Aysegul Nalca, Diane L. Bolton, Margaret LM Pitt).

## Abstract

The COVID-19 pandemic has had a staggering impact on social, economic, and public health systems worldwide. Vaccine development and mobilization against SARS-CoV-2 (the etiologic agent of COVID-19) has been rapid. However, novel strategies are still necessary to slow the pandemic, and this includes new approaches to vaccine development and/or delivery, which improve vaccination compliance and demonstrate efficacy against emerging variants. Here we report on the immunogenicity and efficacy of a SARS-CoV-2 vaccine comprised of stabilized, pre-fusion Spike protein trimers displayed on a ferritin nanoparticle (SpFN) adjuvanted with either conventional aluminum hydroxide or the Army Liposomal Formulation QS-21 (ALFQ) in a cynomolgus macaque COVID-19 model. Vaccination resulted in robust cell-mediated and humoral responses and a significant reduction of lung lesions following SARS-CoV-2 infection. The strength of the immune response suggests that dose sparing through reduced or single dosing in primates may be possible with this vaccine. Overall, the data support further evaluation of SpFN as a SARS-CoV-2 protein-based vaccine candidate with attention to fractional dosing and schedule optimization.

## Introduction

Severe acute respiratory syndrome coronavirus 2 (SARS-CoV-2), the etiologic agent of COVID-19, is responsible for millions of deaths worldwide since the start of the pandemic in late 2019. Despite the fact that licensed and emergency use authorization (EUA) vaccines are available, only about 40% of the world’s population is vaccinated. In the United States, where around 60% of individuals are fully vaccinated, vaccination rates have stagnated due to a variety of factors, including concerns about overall safety of the current vaccines and the need for further testing in young children. Continued spread of SARS-CoV-2 among vulnerable populations has contributed to the emergence of variants. The social, economic, and public health impacts of this ongoing pandemic have been staggering, with a return to normal dependent on the identification of novel strategies to slow the spread of the virus.

Second generation vaccines are currently being developed to supplement or replace existing vaccination strategies and increase immunity and vaccine accessibility. One such approach uses *Helicobacter pylori* ferritin nanoparticles as an antigen scaffold to increase immune responses against a relevant antigen, such as the spike (S) glycoprotein. The S glycoprotein of SARS-CoV-2 is found on the surface of the virus and is responsible for binding to the angiotensin converting enzyme 2 (ACE2) receptor on the surface of epithelial cells and facilitating entry of the virus into the cell [1-3]. Due to its surface location and function, S protein also represents a prominent immunogen of SARS-CoV-2 against which strong neutralizing antibody and cell-mediated responses are mounted [3-5]. The vast majority of vaccines against SARS-CoV-2 utilize the S protein, including the licensed products BNT162b (Pfizer-BioNTech) and mRNA-1273 (Moderna), and the EUA products Ad26.COV2.S (Johnson & Johnson/Janssen) and ChAdOx1nCoV-19 (AstraZeneca) [6-15]. We developed a novel vaccine candidate displaying the S protein on the surface of a *Helicobacter pylori* ferritin nanoparticle (SpFN) described previously [16]. This antigen complex was combined with an adjuvant to promote a robust immune response following vaccination. We previously reported the immunogenicity of SpFN adjuvanted with either aluminum hydroxide (AlOH3) or Army Liposomal Formulation QS-21 (ALFQ) in mice, and observed increased humoral and T cell responses with ALFQ [17]. Subsequent vaccine efficacy studies of SpFN-ALFQ demonstrated protection against disease and viral replication in the respiratory tract of Syrian golden hamsters, and against viral replication in rhesus macaques [18, 19].

In the present study, we evaluated the immunogenicity and efficacy of SpFN adjuvanted with either ALFQ or AlOH3 in a cynomolgus macaque (CM) model of COVID-19. Robust humoral and cellular immunological responses were observed for vaccinated animals, with the ALFQ-adjuvanted group generating a significantly higher response compared to the AlOH3 group. Vaccination with either adjuvant resulted in fewer clinical signs of disease compared to controls, including a significant reduction in lesions and virus replication in the lungs. The data support the continued development of the SpFN adjuvanted vaccine for human use.

## Results

### Vaccination with SpFN reduces clinical disease in SARS-CoV-2-infected CM

A summary of the study design can be seen in Fig 1. Twenty-four adult (3-9 years old) CM passed a veterinary health assessment prior to study assignment, and all animals were assessed and determined negative for prior SARS-CoV-2 exposure using three assays described previously [20]: Euroimmun IgG enzyme linked immunosorbent assay (ELISA), plaque-reduction neutralization test (PRNT), and quantitative real-time PCR (RT-qPCR). This study was conducted in two phases: Vaccination Phase (VP) and Challenge Phase (CP). During the VP, animals randomized into three groups containing eight animals each (balanced by weight and gender) were vaccinated on Study Days -56 and -28 by the intramuscular (IM) route with 50 µg of SpFN formulated with either ALFQ (Group 1) or AlOH3 (Group 2) adjuvant. Group 3 animals were injected IM with PBS on Study Days -56 and -28 and served as study controls. Data Sciences International (DSI) M00 telemetry implants were used for real-time monitoring of body temperature during CP, which commenced on Study Day -7 upon movement into animal biosafety level (ABSL)-3. Telemetry devices implanted for CM #5 and #8 (both in Group 1) ulcerated following surgery, requiring removal, thus no telemetry data are available for these two animals. On Study Day 1, the CM were exposed to 2.89×10^7^ plaque forming units (pfu)/4 mL of SARS-CoV-2 (WA-1) by the intratracheal route, and 3.62×10^6^ pfu/0.5 mL by the intranasal route. Cage side observations were conducted daily to assess clinical signs of disease. Physical examinations, and blood and specimen [bronchoalveolar lavage (BAL) and swab] collections, were conducted under anesthesia on Study Days -4, 1, 3, 5, 7, 9, 11, and 15. None of the animals became terminally ill during the course of the study. Half of the animals in each group were randomly selected for euthanasia on Study Day 9, with the other half euthanized on Study Day 15. This euthanasia schedule was chosen to provide the best opportunity for assessing pathological findings, which are largely absent after Study Day 10, while still assuring that any delay in disease progression in vaccinated animals could be realized [21].

**Fig. 1.**
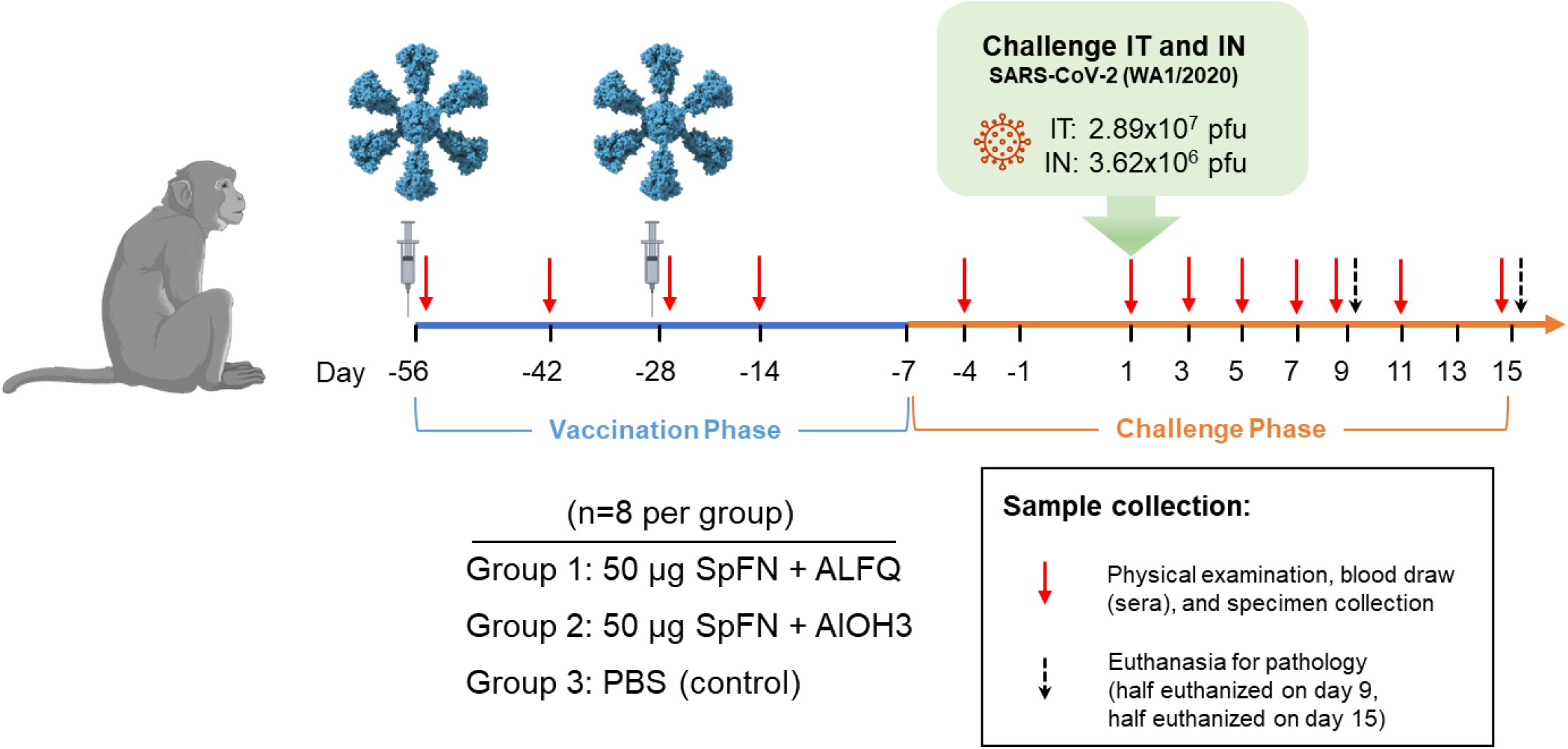
Study Design. Vaccinations occurred on Study Days -56 and -28 as indicated by the syringes in the diagram. Animals were challenged with SARS-CoV-2 (WA-1) on Study Day 1 as indicated. Days of physical examination and blood/specimen collection are indicated by red arrows, and euthanasia days are indicated by black arrows.

Clinical findings are summarized by study group in S1 Table. Disease signs were typically noted between Study Days 2-9. Clinical signs of disease for animals in Group 3 (controls) were similar to those described previously for this model [21-23]. Fever (Fig 2) was the earliest and most consistent finding, was measured for 6/8 animals in this group, was noted between Study Days 2-3, and peaked on Study Day 2. Hyperpyrexia, or an elevation of body temperature greater than 3°C above baseline, was measured for three control animals in Group 3, and all animals in this group had a significantly elevated body temperature (greater than 3 standard deviations above baseline for a duration of greater than 2 hours) on one or more days post-challenge. Although fever and/or significantly elevated body temperatures were noted for 4/8 animals in Group 1 (SpFN + ALFQ), and 1/8 animals in Group 2 (SpFN + AlOH3), the magnitude of the elevation was noticeably less compared to controls (Fig 2). In addition, fever-hours (fever-h), which are the sum of the significant temperature elevation values and give an indication of the intensity of the fever by crudely calculating the area under the curve, were significantly greater for Group 3 control animals compared to animals in the other groups (Fig 2).

**Fig. 2.**
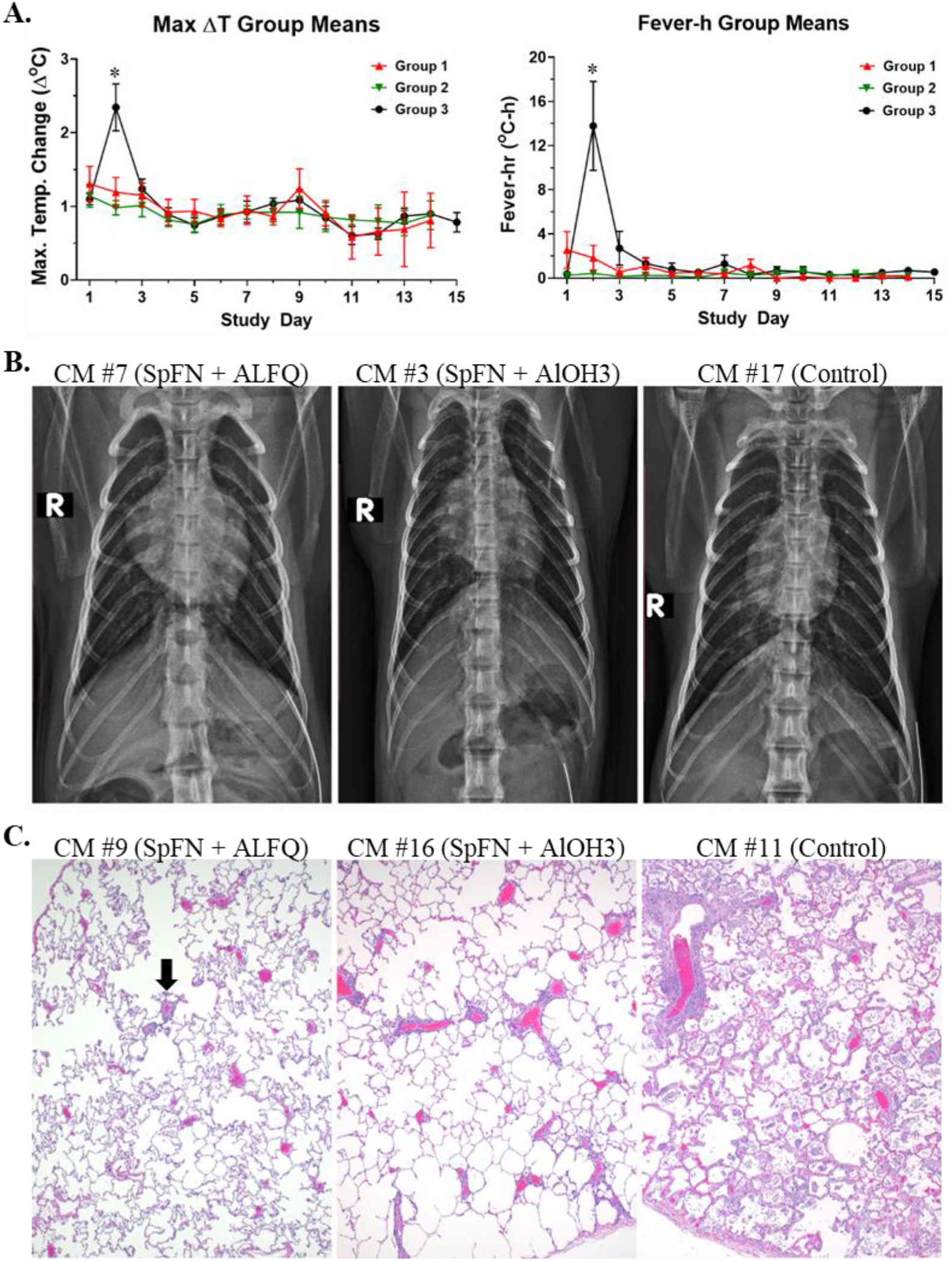
Disease characteristics following SARS-CoV-2 challenge of SpFN-vaccinated CM. A. DSI M00 telemetry devices were used to collect body temperature data. The panel on the left shows the maximum change (Δ) in temperature (^°^C) for the 24-hour daily time period by study group. The panel on the right shows the fever-h measurements by study group, which are the sum of the significant temperature elevation values, and give an indication of the intensity of the fever by crudely calculating the area under the curve. Group 1 (red symbols and lines) = SpFN + ALFQ; Group 2 (green symbols and lines) = SpFN + AlOH3; Group 3 (black symbols and lines) = controls. Errors bars represent the standard deviation for a study group on a particular study day. Significant differences (p<0.05) between Groups 1 and 2, as determined by Wilcoxon rank-sum test, are indicated with a star (*). B. Radiographic findings are shown. Left panel: CM #7 (Group 1, SpFN + ALFQ), Study Day 3, potential very mild opacity increase in the right middle lung lobe. Middle panel: CM #3 (Group 2, SpFN + AlOH3), Study Day 5, moderate infiltrates present in the right middle and caudal lobes, with partial obfuscation of the cardiac silhouette on the right. Right panel: CM #17 (Group 3, control), Study Day 5, partial obfuscation of the right cardiac silhouette with worsening of infiltrate and opacity of middle and caudal lobes bilaterally (R>L); infiltrate also present on the left side. C. Histopathology findings in lung tissue are shown. Left panel: Left cranial lung lobe, CM #9 (Group 1, SpFN + ALFQ), 10X magnification of a peripheral section demonstrating a small amount of perivascular inflammation (black arrow). Middle panel: Left caudal lung lobe, CM #16 (Group 2, SpFN + AlOH3), 10X magnification of a peripheral section demonstrating more perivascular inflammation. Right panel: Right caudal lung lobe, CM #11 (Group 3, control), 10X magnification of a central section demonstrating significantly more inflammation that expands into alveolar spaces, septa and perivascular areas than in the previous images from animals in Groups 1 and 2.

Tachycardia (heart rate greater than or equal to 20 beats per minute above baseline) was a common finding for the majority of animals on this study, regardless of group (S1 Table). Reduced biscuit/enrichment consumption and/or anorexia (absence of biscuit and enrichment consumption for one or more days, or an absence of biscuit consumption for 3 or more consecutive days) appeared to be slightly more common for animals in Groups 2 (SpFN + AlOH3) and 3 (controls), but the difference in the number of animals exhibiting this sign was not significantly different compared to Group 1 (SpFN + ALFQ). In contrast, piloerection was noted for a significantly greater number of animals in Groups 2 and 3 compared to Group 1. Other less common findings (noted for less than or equal to 3 animals in a group) included cough, an abdominal component to breathing, lymphadenopathy (not noted for SpFN + AlOH3 animals), decreased skin turgor (not noted for SpFN + ALFQ animals), not fully formed or liquid stool (not noted for SpFN + ALFQ animals), diurnal rhythm disruptions, and mild hypoxia.

Radiographs were performed on all animals on this study, and the findings are summarized in S1 Table and Fig 2. Overall, disease in the lungs was more pronounced for control animals compared to vaccinated animals, and the lesions noted for Group 2 (SpFN + AlOH3) animals were more common and severe than those noted for Group 1 (SpFN + ALFQ) animals. Although 4/8 animals in Group 1 had evidence of increased lung opacity between Study Days 3-5, this finding was very mild, limited to one lung lobe, and resolved between Study Days 5-7. Furthermore, for CM #8 and #21 of Group 1, this finding was evaluated as a potential positional or rotational artifact. In addition to opacity increases, Group 2 (SpFN + AlOH3) animals exhibited the added finding of infiltrates affecting one or more lobes. For CM #3 in Group 2, lung lesions were extensive enough to cause a partial obfuscation of the cardiac silhouette on the right side on Study Day 5. For Group 2 animals, lesions were present as early as Study Day 3, with resolution between Study Days 7-9. Opacity increases and the presence of infiltrates often affecting multiple lobes were characteristic of the lungs for animals in Group 3 (controls). For CM #13 in Group 3, blurring of right and left cardiac borders was observed on Study Day 5. For CM #15, the right lateral bronchial tree was more apparent compared to baseline images on Study Day 3. For CM #17, partial obfuscation of the right cardiac silhouette was noted on Study Day 5. There also appeared to be a delay in resolution of lesions for animals in Group 3 (controls) compared to Group 2 (SpFN + AlOH3). Progression of lung disease was noted as late as Study Day 7 for CM #13 and #19 in Group 3, and incomplete resolution of lung lesions was noted as late as Study Day 9 for CM #13 (end-of-study for this animal), CM #15, CM #17 (end-of-study for this animal), and CM #19 in Group 3. For CM #19, improved but incomplete resolution was observed on Study Day 15.

At necropsy, fibrosis was noted for 2/8 animals in Group 1 (SpFN + ALFQ), and 1/8 animals in Group 3 (control) had fibrin tags extending from the right caudal lung lobe to the pleura of the thoracic wall. Otherwise, gross findings at necropsy were limited, with only typical post-mortem changes found. While histological lesions across all groups were generally mild, vaccination with SpFN, regardless of adjuvant, appeared to result in less inflammation and very little to no damage to alveolar septa. The character and distribution of inflammation in the lungs and nasal turbinates observed in this study was consistent with that previously reported for CM exposed to SARS-CoV-2 [20, 21] (S2 Appendix), and the findings in the lungs were also consistent with radiographic findings for this tissue. Inflammation was more frequent and prominent for control animals (Fig 2 and S2 Appendix). Reparative lesions, such as alveolar fibrosis and type II pneumocyte hyperplasia, were largely observed only for Group 3 (control) animals, with the exception of a single animal in Group 2 (SpFN + AlOH3). Viral antigen detected by immunohistochemistry was only present in lung sections from control animals.

Given their proximity to the lungs, the tracheobronchial and axillary lymph nodes represent prominent draining lymph nodes for this SARS-CoV-2 model. Lymphoid hyperplasia of the tracheobronchial lymph node was common (greater than 50% incidence) across all groups (S1 Appendix). The axillary lymph node also exhibited lymphoid hyperplasia, but at a lower rate in all groups (S1 Appendix). These findings are attributed to stimulation of the adaptive immune response to either vaccine administration and/or exposure to SARS-CoV-2.

### Clinical pathology analyses

Complete blood counts and clinical chemistries (Fig 3, S2 Table, and S3 Table) were performed on whole blood and serum, respectively. Clinical pathology changes during VP were largely unremarkable (S2 Table), with the exception of creatine kinase which was elevated above baseline for the majority of animals on at least one or more VP study day. The reason for this elevation is unknown but was most likely related to stress or possibly inflammation in muscle tissue following injections.

**Fig. 3.**
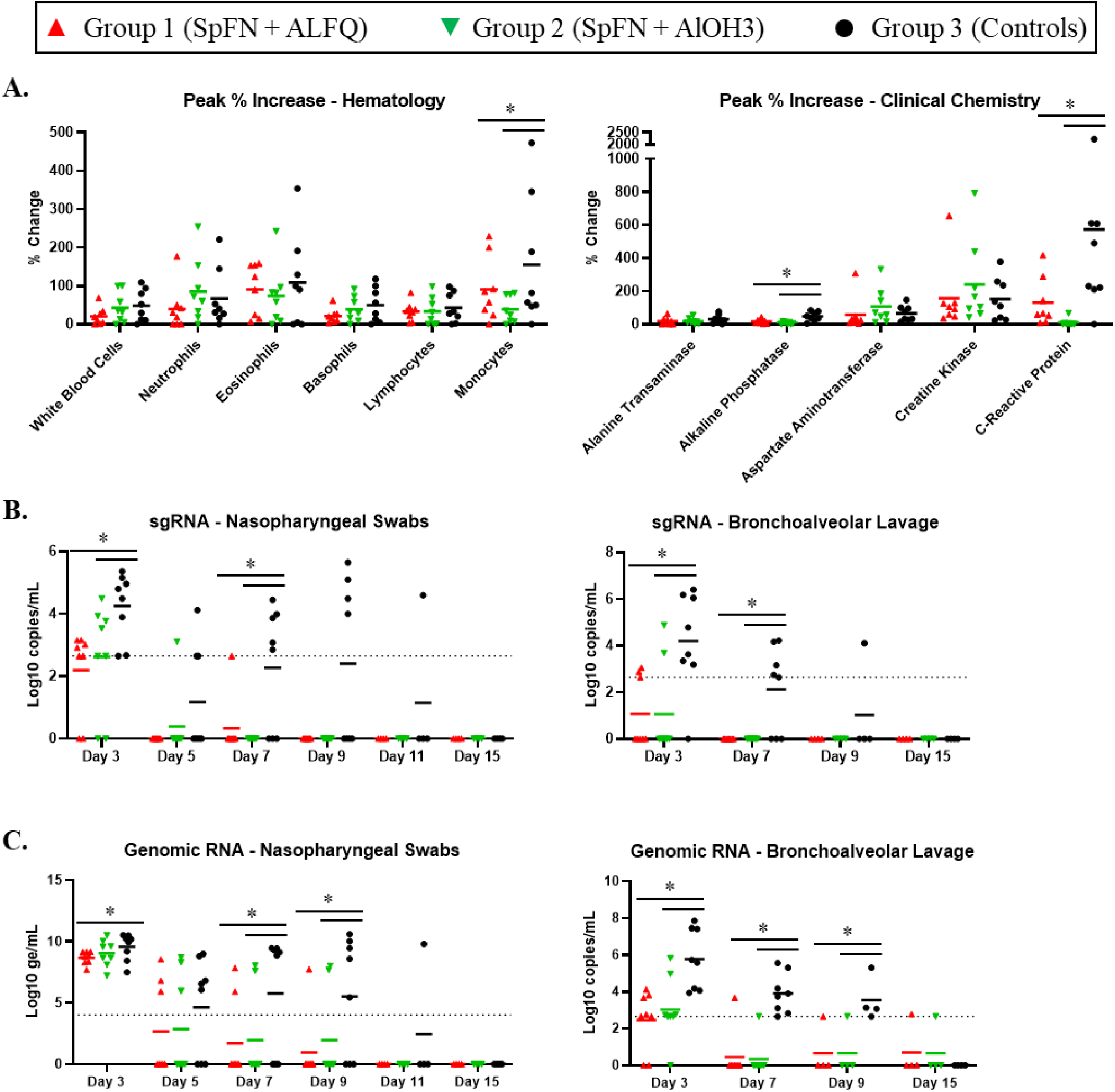
Clinical pathology and respiratory tract-associated viral RNA levels following SARS-CoV-2 challenge of SpFN-vaccinated CM. Data are presented by group. Group 1 (red symbols and lines) = SpFN + ALFQ; Group 2 (green symbols and lines) = SpFN + AlOH3; Group 3 (black symbols and lines) = controls. Significant differences (p<0.05) between control and vaccinated, as determined by Wilcoxon rank-sum test, are indicated with a star (*). Horizontal bars represent the mean value. A. Clinical pathology. Measurements are shown as percent change from baseline (average of values from Study Day -4 and 1 for each animal) for peak values for each analyte. The panel on the left shows hematology data, and the panel on the right shows clinical chemistry data. B. Subgenomic RNA in nasopharyngeal (NP) swabs (left panel) and BAL (right panel). Data are shown as Log_10_ copies/mL. The dotted lines demarcate assay lower limit of linear performance range (Log_10_ = 2.65, corresponding to 450 copies/mL). C. SARS-CoV-2 RT-qPCR to detect genomic RNA was performed on NP swabs (left panel) and BAL (right panel). Data are shown as Log_10_ genomic equivalents (ge)/mL. The dotted lines demarcate assay lower limit of linear performance range for BAL (Log_10_ = 2.65) or the lower limit of quantification for NP swabs (Log_10_ = 4.00).

Most changes in clinical pathology parameters during CP were consistent with those noted previously [20, 21], and can be attributed to an inflammatory response indicative of ongoing virus infection. This included increases in one or more leukocyte parameters, as well as increases in creatine kinase and/or C-reactive protein (Fig 3 and S3 Table). In general, any changes observed were similar between groups with two exceptions: C-reactive protein and monocytes. In both cases, the change from baseline was significantly greater for control animals compared to vaccinated animals.

Although increases in one or more liver-related enzyme activities were noted across groups, the changes were minimal and likely not biologically meaningful. This is further supported by the histopathologic findings which indicated that liver disease was largely absent at the time of necropsy (S1 Appendix).

### Viral replication in the respiratory tract is reduced for SpFN-vaccinated CM

To assess the impact of vaccination on viral replication in the respiratory tract, SARS-CoV-2 subgenomic RNA (sgRNA) was assessed in BAL and nasopharyngeal (NP) swabs by quantitative reverse transcription polymerase chain reaction (RT-qPCR, Fig 3). Two days following challenge (Study Day 3), sgRNA in control animals averaged 10^4^ copies/mL in both BAL and NP swabs, with 7/8 animals exhibiting robust viral replication in the BAL, and all animals exhibiting replication in the nasopharyngeal tract. Peak (Study Day 3) BAL and NP swab titers were significantly higher for control animals compared to vaccinated animals. Among vaccinated animals, sgRNA was below the limit of detection (LOD) in BAL of 5/8 and 6/8 animals in the ALFQ and AlOH3 groups, respectively. By Study Day 7 (6 days post-challenge), 5/8 controls had detectable replication in the lungs, while all vaccinated animals had undetectable BAL sgRNA. BAL sgRNA was resolved for all but one control animal by Study Day 9 (8 days post-challenge). Similar trends were observed when the NP swab material was assessed, with sgRNA present in NP swabs of 5/8 controls on Study Day 7, but absent in all AlOH3 animals and all but one ALFQ animal. SgRNA persisted in NP swabs of 4/8 control animals through Study Day 9, while all vaccinated animals were below the LOD at that time point.

Genomic RNA was also assessed on BAL and NP swab samples by RT-qPCR (Fig 3). In general, trends were similar to those seen for sgRNA. BAL genomic RNA levels for vaccinated animals were significantly lower than control animals on Study Day 3 (2 days post-challenge). Unlike control animals, for which viral RNA was still detected on Study Day 9 (8 days post-challenge), viral RNA in BAL was detected beyond Study Day 3 for only one animal in each vaccine group. All animals had quantifiable viral RNA in NP swabs by Study Day 3 (2 days post-challenge), with mean values of 9.57, 8.68, and 9.05 Log_10_ genomic equivalents (ge)/mL for controls, ALFQ, and AlOH3 groups, respectively. These peak levels were significantly higher among controls relative to SpFN + ALFQ vaccinated animals. By Study Day 5 (4 days post-challenge), less than half of the vaccinated animals in Groups 1 (SpFN + ALFQ) and 2 (SpFN + AlOH3) had detectable viral RNA in swabs, and by Study Day 7 (6 days post-challenge) viral RNA levels were below detection for nearly all of the animals in these two groups. Whereas viral RNA in swabs persisted through Study Day 9 (8 days post-challenge) for the majority of control animals in Group 3.

### Vaccination with SpFN elicits a strong SARS-CoV-2-specific antibody response

The overall IgG and IgA response to SARS-CoV-2 was assessed using the Euroimmun ELISA on samples collected during the VP and the CP, and characterization of the antigen-specific IgG and IgM response was performed by Magpix multiplex immunoassay (Fig 4). An IgG response was detected for vaccinated animals two weeks following initial vaccination (on Study Day -42), with ALFQ group having a significantly stronger response than the AlOH3 group. The second (booster) vaccination on Study Day -28 enhanced the antibody response detected by Study Day -14 for both groups, which had similar IgG levels at all subsequent time points. An anamnestic response for total IgG was not seen following challenge. Characterization of the antigen-specific IgG response by Magpix for vaccinated animals revealed the greatest binding to S1 both pre- and post-challenge. In addition, the level of S1 binding antibodies increased following challenge, demonstrating an anamnestic response to this antigen. Although detectable, the response to full spike was much lower compared to S1, and comparable in magnitude to the response mounted against RBD.

**Fig. 4.**
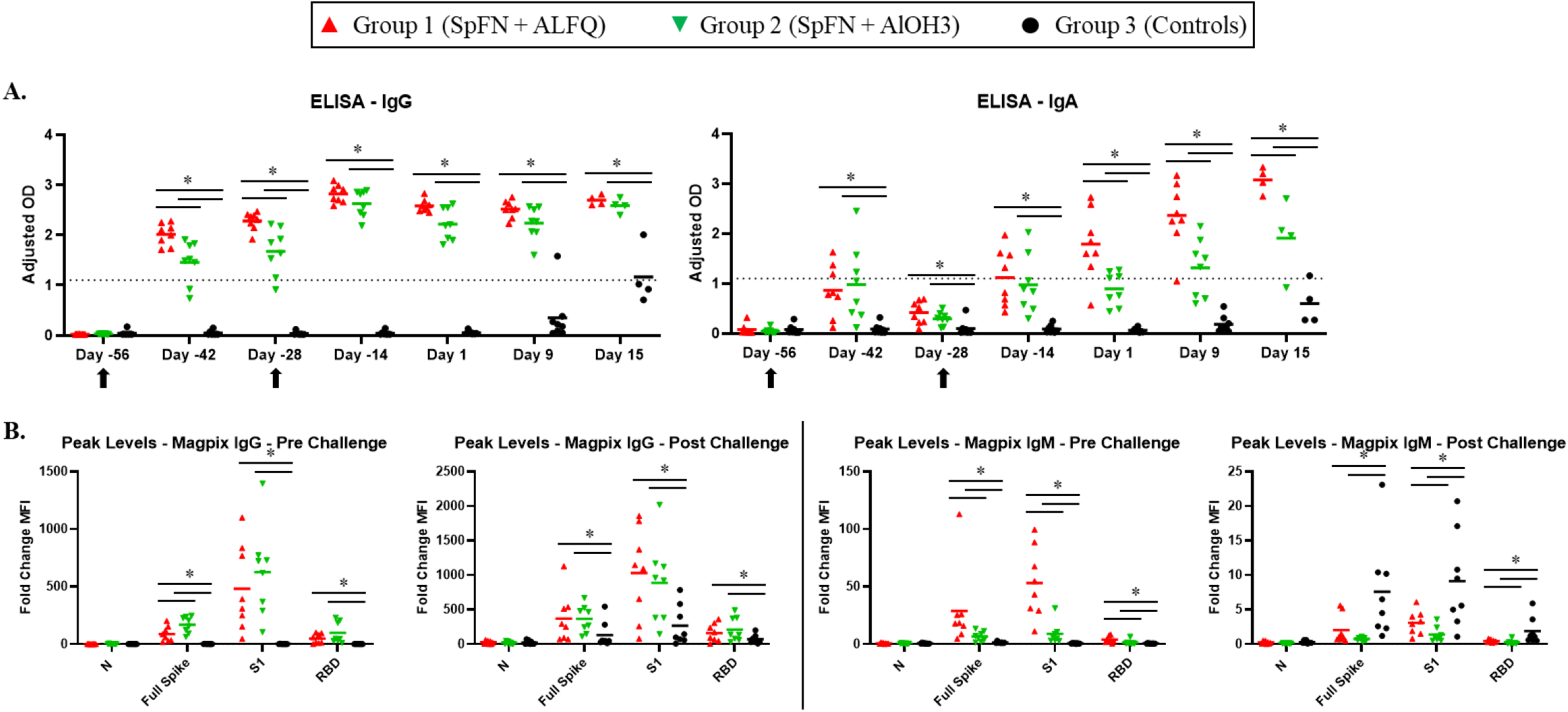
Characterization of the total S1-specific antibody response using ELISA, and antigen-specific antibody response by MagPix. Data are presented by group. Group 1 (red symbols and lines) = SpFN + ALFQ; Group 2 (green symbols and lines) = SpFN + AlOH3; Group 3 (black symbols and lines) = controls. Significant differences (p<0.05) between control and vaccinated, as determined by Wilcoxon rank-sum test, are indicated with a star (*). A. Serum samples were assessed using the Euroimmun SARS-CoV-2 S1 IgG or IgA ELISA kit. The dotted line represents the assay cutoff, above which a sample is considered above background noise (i.e. positive). The total S1 IgG response is shown in the left panel, and the total S1 IgA response is shown in the right panel. Black arrows indicate the days of vaccination. OD = optical density (Adjusted OD is the OD resulting following removal of background noise). B. The IgG (left panels) and IgM (right panels) responses to the indicated antigens were measured by a Magpix immunoassay.

A minimal total IgA response (Fig 4A) was mounted following initial vaccination for vaccinated animals, but by Study Day -28 levels had dropped considerably. Following the second vaccination on Study Day -28, a second IgA response was noted, and levels continued to rise for Group 1 (SpFN + ALFQ) animals over the remainder of the study. Unlike Group 1, levels of IgA for Group 2 (SpFN + AlOH3) plateaued briefly after Study Day -14. For this group, a slight increase was noted by Study Day 9, with an even greater elevation on Study Day 15. The total IgA response was significantly higher for the ALFQ-adjuvanted group on Study Days 1, 9, and 15 compared to the AlOH3-adjuvanted group.

The total IgM response was not measured by ELISA on this study, as Euroimmun kits were only commercially available for IgA and IgG. However, the antigen-specific IgM response was evaluated using the Magpix assay (Fig 4B). Similar to the IgG data, animals in Group 1 (SpFN + ALFQ) developed an IgM response following vaccination to the S1 subunit and, to a much lesser degree, full spike; an enhancement of this response was not seen for post-challenge samples. Contrastingly, the IgM response for Group 2 (SpFN + AlOH3) animals was largely unremarkable.

IgG and IgA antibody responses for control animals (Group 3) were not noted before Study Day 9, and levels were significantly lower than vaccinated animals at all time points assessed (Fig 4A). However, the IgM response to S and full spike was significantly higher for Group 3 control animals in post-challenge samples compared to vaccinated animals (Fig 4B).

Total serum binding responses to SARS-CoV-2 WA-1 and five VoC receptor-binding domain (RBD) molecules were analyzed using biolayer interferometry (BLI), with responses greatest against WA-1, Alpha and Delta VoC (Fig 5). This was consistent for both Groups 1 and 2, with the ALFQ-adjuvanted Group 1 showing significantly higher binding responses than the AlOH3-adjuvanted Group 2 against each VoC.

**Fig. 5.**
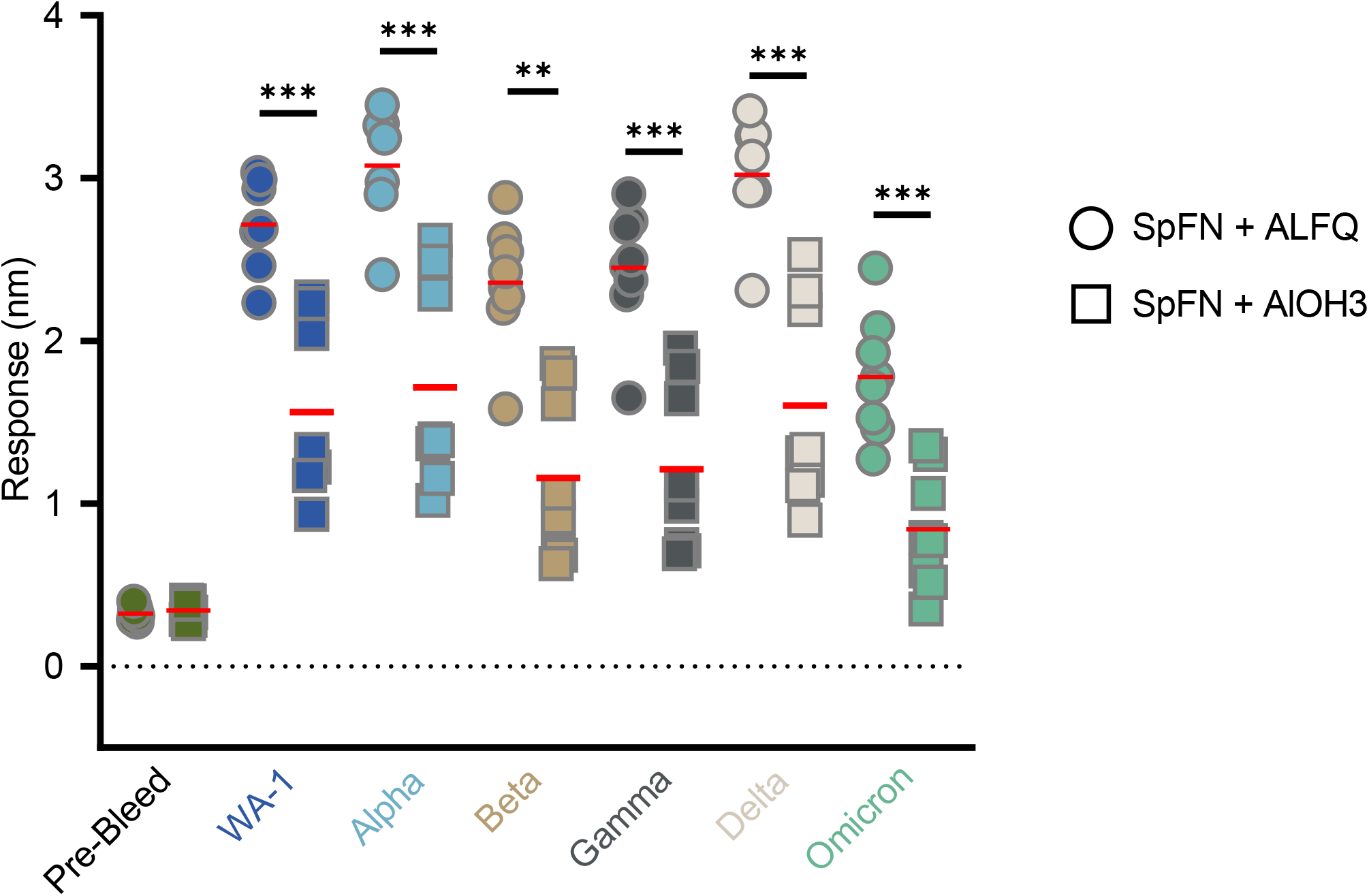
Total serum RBD binding response following SpFN vaccination in CM. Serum collected from all animals on Study Day -56 or -14 was analyzed by BLI for binding to SARS-CoV-2 WA-1 and VoC RBD molecules. Data are presented by group. Group 1 (Circles) = SpFN + ALFQ; Group 2 (Squares) = SpFN + AlOH3. The horizontal bars represent the mean for each group. Groups were compared using unpaired non-parametric two-tailed Mann-Whitney test, where * is p<0.05, ** is p<0.01, and *** is p<0.001.

SARS-CoV-2-specific binding IgG antibody responses, and the ability of S-specific binding antibodies to inhibit S or RBD binding to the ACE-2 receptor, were measured using a Meso Scale Discovery (MSD) immunoassay (Fig 6). Vaccination with SpFN + ALFQ was associated with significantly higher levels of S and RBD-specific IgG antibodies against WA-1 and the Alpha, Beta, and Gamma variants of SARS-CoV-2 compared to vaccination with SpFN + AlOH3. Similarly, SpFN + ALFQ induced significantly higher ACE2 inhibitory antibodies that blocked ACE2 binding to S and RBD of WA-1 and the Alpha, Beta, and Gamma variants compared to SpFN + AlOH3.

**Fig. 6.**
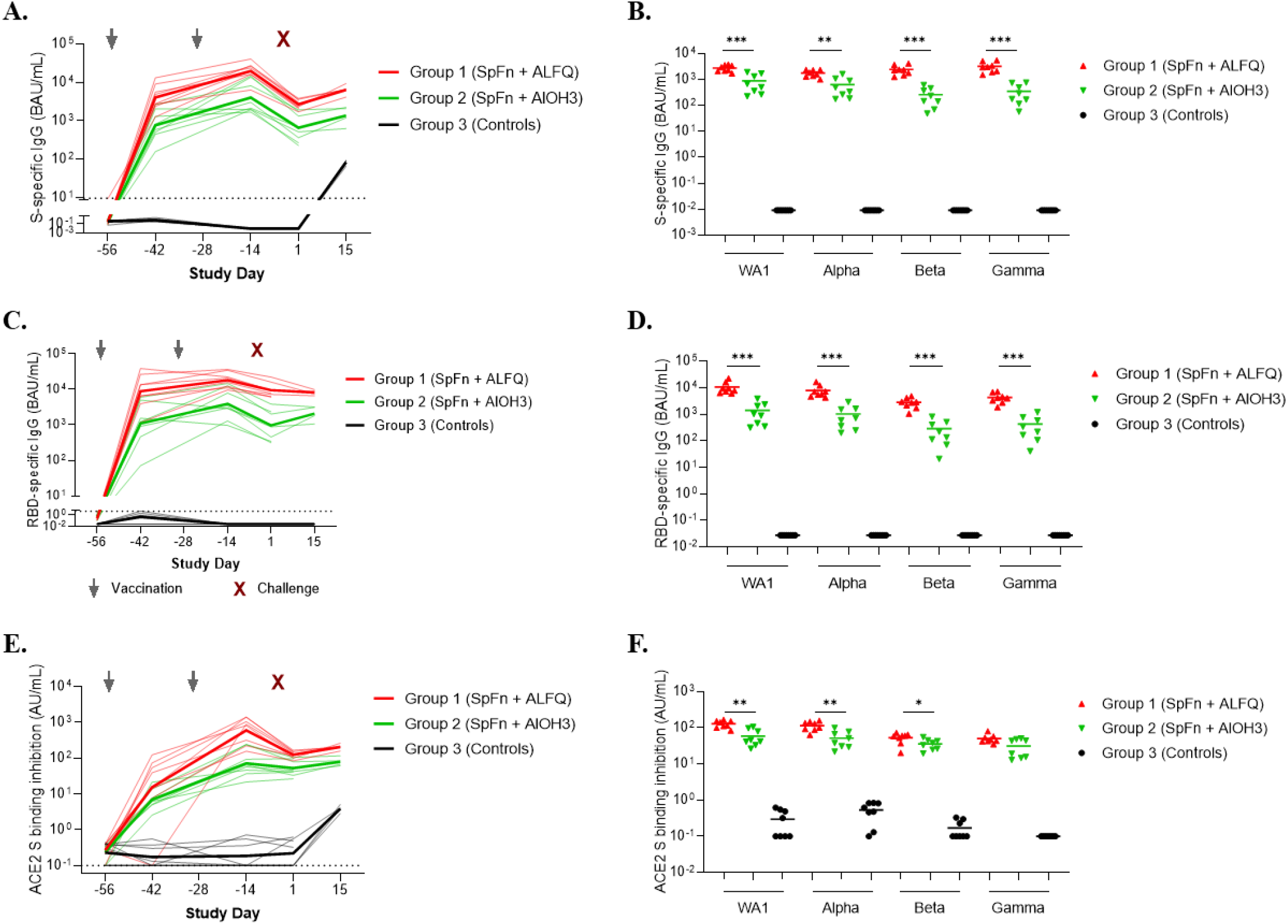
Adjuvanted SpFN vaccine-elicited Spike and RBD binding and ACE-2 inhibitory responses to SARS-CoV-2 assessed by MSD immunoassay. Humoral responses were measured by MSD immunoassay. Data are presented by group. Group 1 (red symbols and lines) = SpFN + ALFQ; Group 2 (green symbols and lines) = SpFN + AlOH3; Group 3 (black symbols and lines) = controls. Dotted lines represent geometric mean of pre-immune values plus five standard deviations, considered the threshold for positive responses. A. Serum SARS-CoV-2 (WA-1) S-specific IgG responses are depicted as binding antibody units/mL (BAU/mL), using the WHO International Standard and a conversion factor of 0.009. Thick lines indicate geometric means for each group and thin lines represent individual animals. B. Serum IgG binding antibody responses to S antigens from SARS-CoV-2 WA-1, Alpha, Beta, and Gamma on Study Day 1. Significant differences between ALFQ and AlOH3-adjuvanted SpFN are indicated (*, p<0.05; **, p<0.01; ***, p<0.001; Mann-Whitney test). C. WA-1 RBD-specific serum IgG binding responses are depicted longitudinally as BAU/mL (conversion factor 0.027). D. Serum IgG binding antibody responses against RBD antigens from WA-1, Alpha, Beta, and Gamma. Significant differences between ALFQ and AlOH3-adjuvanted SpFN are indicated. E. Serum inhibition of SARS-CoV-2 (WA-1) S binding to angiotensin-converting enzyme 2 (ACE2) reported as arbitrary units (AU/mL). F. Serum inhibition of ACE-2 binding to SARS-CoV-2 S proteins from WA-1, Alpha, Beta, and Gamma on Study Day 1. Significant differences between ALFQ and AlOH3-adjuvanted SpFN are indicated.

The neutralizing antibody response, assessed by plaque reduction neutralization test (PRNT), was significantly higher for Group 1 (SpFN + ALFQ) compared to Group 2 (SpFN + AlOH3) on Study Days -42 and -28 (Fig 7). Following the second (boost) vaccination, a significant difference was still observed between vaccinated groups through Study Day 9. Similar to total IgG by ELISA, an anamnestic response for neutralizing antibodies was not seen following challenge.

**Fig. 7.**
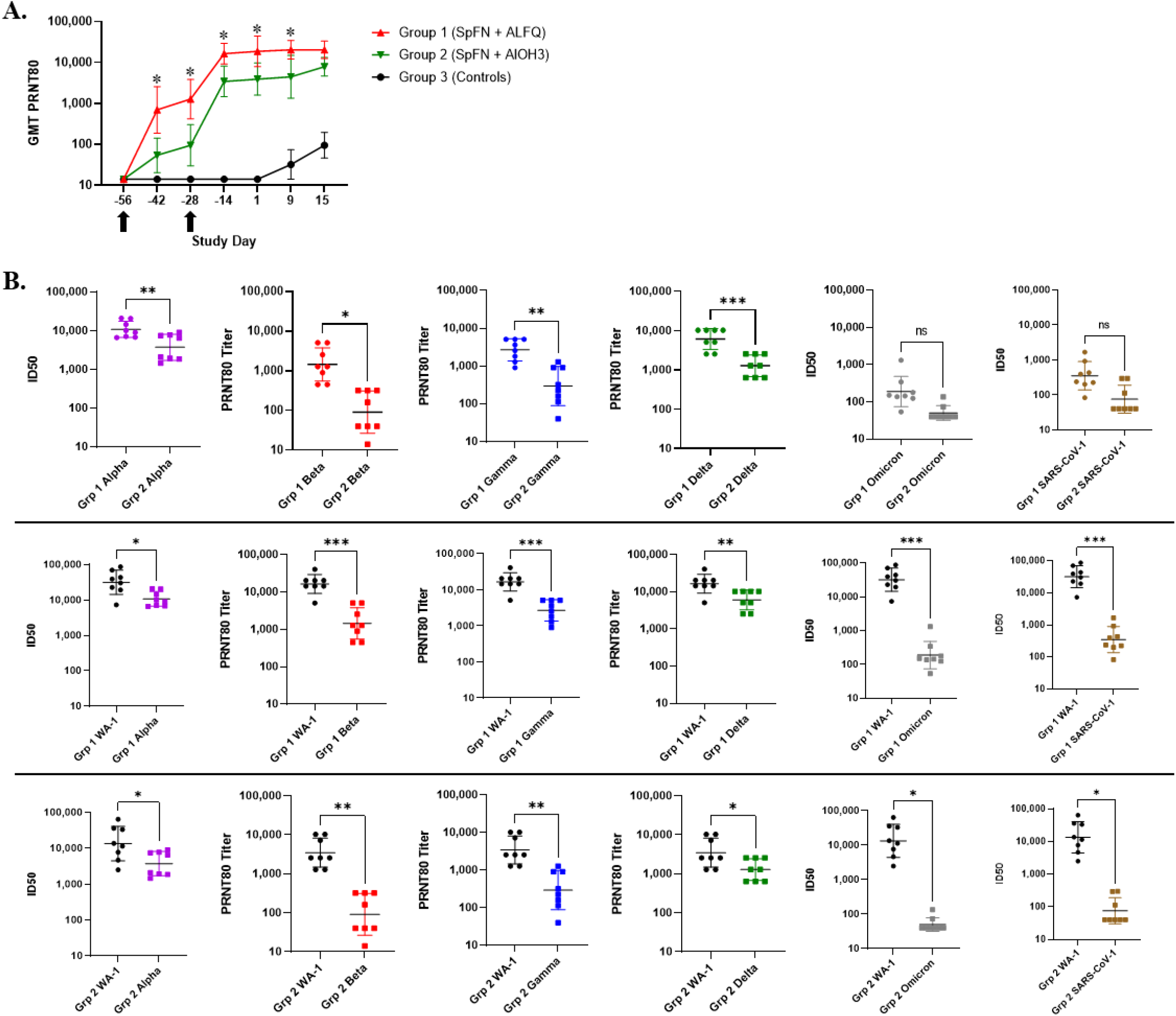
SARS-CoV-2 neutralizing antibody responses following SpFN vaccination in CM. A. The PRNT80 titers to the WA-1 strain of SARS-CoV-2 are shown. Data are presented by group. Group 1 (red symbols and lines) = SpFN + ALFQ; Group 2 (green symbols and lines) = SpFN + AlOH3; Group 3 (black symbols and lines) = controls. Significant differences (p<0.05) between Groups 1 and 2, as determined by Wilcoxon rank-sum test, are indicated with a star (*) (Note: Comparison of Group 3 to Groups 1 and 2 revealed statistical significance by Wilcoxon rank-sum test at all post-vaccination time points). Black arrows indicate the days of vaccination. GMT = geometric mean neutralization titer. Error bars represent the geometric standard deviation. B. The 50% infectious dose (ID50) geometric mean (pseudovirion assay, Alpha and Omicron variants and SARS-CoV-1) and PRNT80 GMTs (live virus assay, Beta, Gamma, and Delta variants) against variants of concern (VoC) are shown. Error bars represent the standard deviation. Statistical relevance was determined using unpaired t tests, where * is p<0.05, ** is p<0.01, and *** is p<0.001. In the top panels, Group 1 (SpFN + ALFQ) and Group 2 (SpFN + AlOH3) titers are compared for each VoC. In the middle panels, titers for Group 1 against WA-1 are compared to those measured for VoC. In the bottom panels, titers for Group 2 against WA-1 are compared to those measured for VoC.

Live virus PRNTs and pseudovirion neutralization assays were also performed on serum samples to look at neutralization of five SARS-CoV-2 variants of concern (VoC)- Alpha (or B1.1.7, first identified in United Kingdom) (pseudovirion assay only), Beta (or B1.351, first identified in the Republic of South Africa), Gamma (or P.1, first identified in Brazil) (live virus assay only), Delta (or B1.617.2, first identified in India) (live virus assay only), and Omicron (or B1.1.529, first identified in southern Africa) (Fig 7). Although there was significant reduction in neutralizing activity against the VoC compared to assay-matched WA-1, the Group 1 (SpFN + ALFQ) response to Alpha, Beta, Gamma, and Delta was still relatively strong. This contrasts with the responses measured for Group 2 (SpFN + AlOH3) which were not only significantly lower than to WA-1, but were also significantly lower for Alpha, Beta, Gamma, and Delta compared to Group 1. Only a low-level response against Omicron was measured for both vaccine groups.

Pseudovirion assays were also performed to look at cross-neutralization of SARS-CoV-1 following vaccination (Fig 7B). Although the response to SARS-CoV-1 was significantly lower than the response to WA-1 and VoC for vaccinated groups, a low-level response was still measured. This data suggested that SpFN adjuvanted with either ALFQ or AlOH3 might have a minimal cross-protective capability against SARS-CoV-1.

Non-neutralizing antibody effector functions are associated with vaccine-mediated protection against other viruses [24, 25], and may also be important for protection against SARS-CoV-2 [9, 26]. Two weeks following the initial vaccination (Study Day -42), antibody dependent complement deposition (ADCD) and antibody dependent cellular phagocytosis (ADCP) levels were significantly higher for Group 1 (SpFN + ALFQ) animals compared to Group 2 (SpFN + AlOH3) animals (Fig 8). Both groups had strong ADCD and ADCP levels that were similar in magnitude two weeks following the boost vaccination (Study Day -14).

**Fig. 8.**
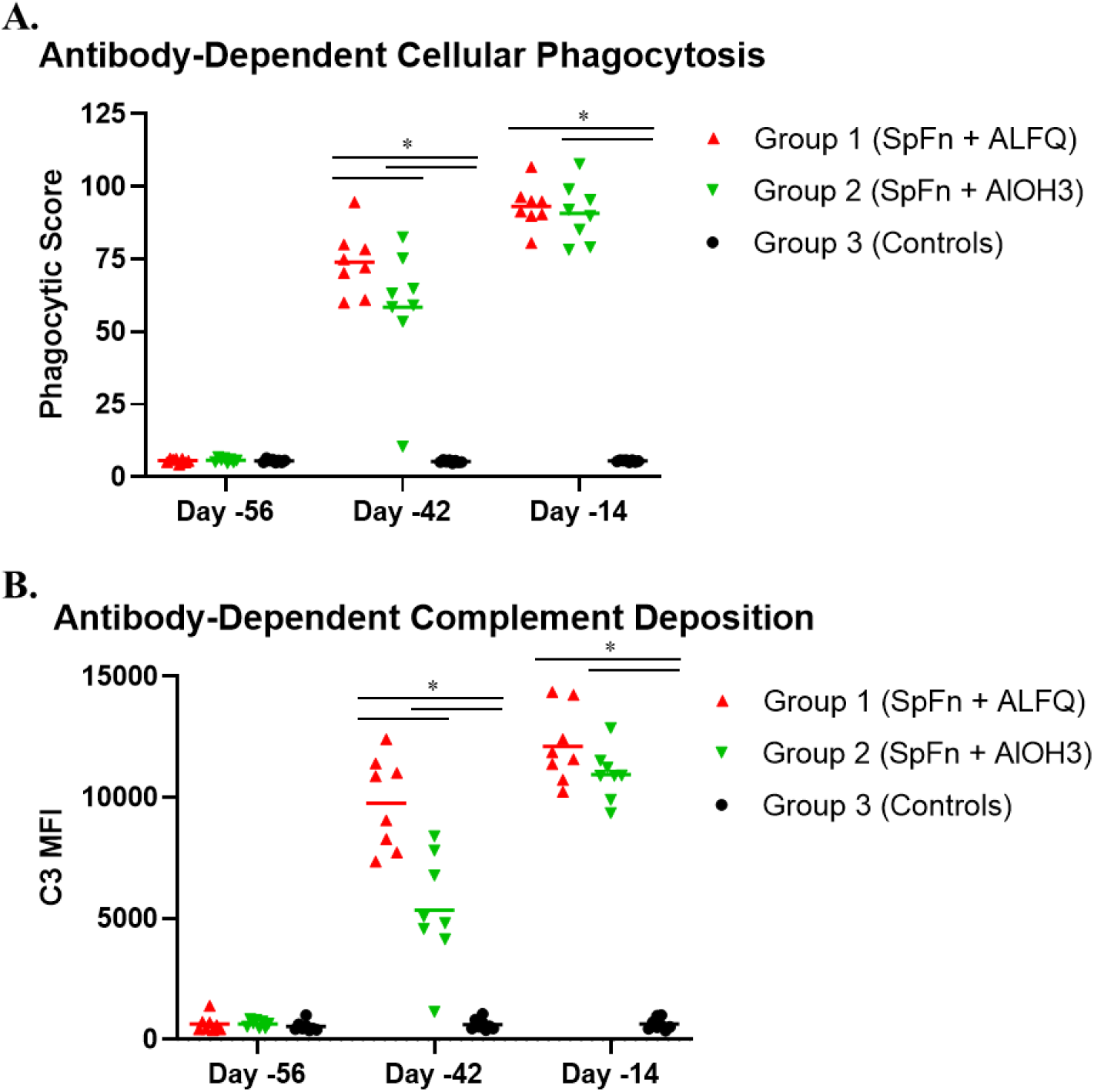
Non-neutralizing antibody effector functions following SpFN vaccination in CM. Data are presented by group. Group 1 (red symbols and lines) = SpFN + ALFQ; Group 2 (green symbols and lines) = SpFN + AlOH3; Group 3 (black symbols and lines) = controls. Significant differences (p<0.05) between control and vaccinated, as determined by Wilcoxon rank-sum test, are indicated with a star (*). A. ADCP. B. ADCD. MFI = mean fluorescence intensity.

### SARS-CoV-2-specific T cell responses elicited by vaccination with SpFN

SARS-CoV-2-specific T cell immunity contributes to control and resolution of infection in addition to supporting antibody responses [27-29]. S-specific T cells were characterized in peripheral blood mononuclear cells (PBMCs) by *in vitro* peptide pool stimulation and intracellular cytokine staining (ICS) using a 19-color multi-parameter flow cytometry (Fig 9, S1 Fig, S2 Fig). Two weeks following the boost, a significant Th1 (TNF-α, IFN-*γ*, IL-2) CD4+ T cell response was elicited by adjuvanted SpFN in all but one animal, ranging from 0.8-8% of memory CD4+ T cells in the SpFN + ALFQ group (Group 1) and 0.5-1.9% in the SpFN + AlOH3 group (Group 2). Th2 responses (IL-4 and IL-13) and CD8+ T cell responses were low or undetectable. To evaluate additional CD4+ T cell functions important for B cell development and antibody responses, S-specific CD4+ T cells expressing IL-21 and CD40L were also measured. Seven of eight SpFN + ALFQ and five of eight SpFN + AlOH3 vaccinated animals developed IL-21 responses following boost (Fig 9C), with mean responses of 0.5% and 0.08% of memory CD4+ T cells, respectively. S-specific CD40L+ CD4+ T cell responses were similar in magnitude and kinetics to the Th1 response, again with notable post-boost peak responses of 1.0-8.8% elicited by SpFN + ALFQ and 1.1-2.1% in six of eight animals that received SpFN + AlOH3 (Fig 9D).

**Fig. 9.**
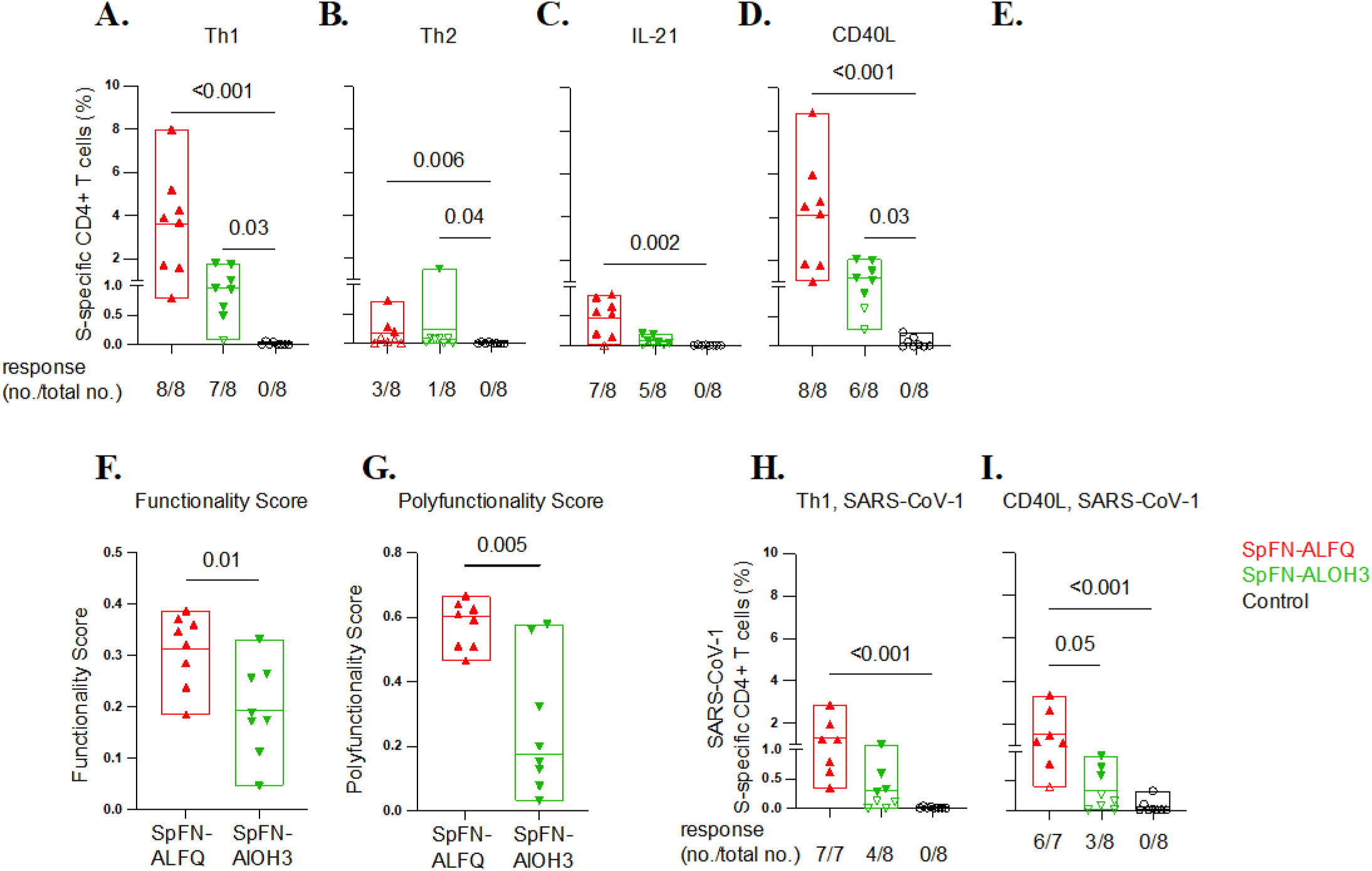
SARS-CoV-2 and SARS-CoV-1 S-specific CD4+ T cell responses elicited by adjuvanted SpFN vaccination. A-D. T cell responses were assessed by SARS-CoV-2 spike peptide pool stimulation and intracellular cytokine staining of PBMC collected at Study Day -14 (2 weeks post-boost). S-specific memory CD4+ T cells were defined by expression of: A. Th1 cytokines (IFN-*γ*, TNF-*α* and IL-2), B. Th2 cytokines (IL-4 and IL-13), C. IL-21 or D. CD40L. Boolean combinations of cytokine-positive memory T cells were summed for Th1 and Th2 response magnitude. E. Pie charts depict polyfunctionality of S-specific Th1 CD4+ T cell responses 2 weeks post-boost for the two adjuvanted SpFN vaccine arms, as assessed by Boolean combination gating of IFN-*γ*, TNF-*α* and IL-2 positive cells. F-G. COMPASS analysis of S-specific CD4+ T cell functionality F. and polyfunctionality G. determined by Th1 cytokine (TNFα, IL2, IFN*γ*), CD40L, and IL-21 expression. H-I. SARS-CoV-1 S-specific CD4+ T cell responses were measured by PBMC stimulation with SARS-CoV-1 spike peptide pools. The frequency of memory CD4+ T cells expressing Th1 cytokines (IFN-*γ*, TNF-*α* and IL-2) H. and CD40L I. is shown. Study groups are depicted as follows: SpFN + ALFQ, red; SpFN + AlOH3, green; controls, black. Box plot horizontal lines indicate the mean; top and bottom reflect the minimum and maximum. The fraction of animals within each group with a positive response following each vaccination is indicated. Significance was assessed using a Kruskal-Wallis test followed by a Dunn’s post-test to compare all study groups, or a Mann-Whitney test to compare the vaccine groups.

The S-specific CD4+ Th1 cell responses were also polyfunctional following the boost (Fig 9E). On average, 25% and 8% of the S-specific cells produced all three Th1 cytokines among SpFN + ALFQ and SpFN + AlOH3 vaccinated animals, respectively, while 50% expressed two of the three cytokines. T cell response functionality was also assessed by COMPASS, a high-dimensional ICS data analysis tool to identify antigen-specific T cell subsets and define their relationship to clinical outcomes [30]. Boolean combinations of cells expressing Th1 cytokines (TNF-α, IL-2, IFN-*γ*), CD40L, and IL-21 were used to calculate functionality and polyfunctionality scores following immunization. S-specific CD4+ T cell functionality and polyfunctional scores were both higher following immunization with SpFN + ALFQ than with SpFN + ALOH3 (Fig 9F-G), consistent with ALFQ modulation of SpFN-elicited T cell responses to greater polyfunctionality.

Given the emergence of SARS-CoV-2 variants of concern that may evade adaptive immune responses elicited by vaccination, cross-reactive T cell responses to the divergent SARS-CoV-1 S protein were assessed using SARS-CoV-1 S peptide pools and ICS as described above. SARS-COV-1 S-specific Th1 CD4+ T cell responses were elicited in all SpFN + ALFQ vaccinated animals (0.3-2.9%), and half of the SpFN + AlOH3 animals (0.3-1%) following the boosting immunization (Fig 9H). Similarly, significant CD40L responses against SARS-CoV-1 S were generated by SpFN + ALFQ prime-boost, with more modest cross-reactive responses with AlOH3 adjuvant (Fig 9I). Overall, these data show that SpFN adjuvanted with ALFQ, and at a lesser extent with AlOH3, induced robust Th1-polarized polyfunctional CD4+ T cells favorable for viral clearance and with critical B cell help capability and cross-reactivity to divergent SARS-CoV-1 sequence.

## Discussion

The development of effective and durable vaccines against SARS-CoV-2 is a critical to addressing the COVID-19 pandemic. Although FDA approved and EUA vaccines are available for use against SARS-CoV-2, vaccination rates have stagnated, falling short of required percentages to assure herd immunity. The considerable size of the high risk and vulnerable unvaccinated population, coupled with the emergence of variants due to sustained circulation in the human population, has allowed for continued rapid spread of SARS-CoV-2. Additional resources, including novel approaches to vaccine development, are critical for a return to normal. Herein, we describe the results of a statistically-balanced nonhuman primate study in which we evaluated the efficacy of SARS-CoV-2 SpFN protein nanoparticle vaccine against a robust and high-titer SARS-CoV-2 respiratory (IT) and mucosal (IN) challenge. Efficacy and/or immunogenicity of this vaccine has been demonstrated in mice, hamsters, and rhesus macaques [17, 19, 31], including efficacy against the Alpha and Beta variants of SARS-CoV-2 demonstrated in hamsters [19]. CM were chosen for the present study as this model reproduces several human disease characteristics and provides two ideal objective and relevant endpoint criteria for efficacy evaluations: fever and viral RNA in nasopharyngeal swabs [20, 21].

Clinical disease findings for virus-only control animals (Group 3) were similar to those previously described [20, 21] and included fever, piloerection, and reduced food consumption. Evidence of disease in the lungs was present both radiographically and histopathologically. SgRNA, which is an indicator of viral replication, was high in BAL and nasopharyngeal swab material, suggesting active viral infection in the lungs and nasopharyngeal tissue.

Vaccination with SpFN did not appear to be associated with any significant adverse reactions or severe findings in CM, consistent with a prior study evaluating SpFN + ALFQ in rhesus macaques [31]. Vaccination with SpFN adjuvanted with either AlOH3 or ALFQ significantly reduced the prevalence, duration, and magnitude of fever, as well as the severity of lesions in the lungs. A significant reduction in the amount of sgRNA present in BAL fluid and nasopharyngeal swab material was also noted, and sgRNA was detectable for far fewer days compared to controls. Similar to screening performed for human patients to confirm cases of COVID-19, nasopharyngeal swab material was assessed by RT-qPCR for the presence of genomic RNA. Although genomic RNA was present at quantifiable levels in the nasopharyngeal swabs of vaccinated animals at early time points (Study Day 3), sgRNA in the same tissue was generally below the LOD. Taken together, these results indicate that the majority of the viral RNA detected in swab material after Study Day 3 for vaccinated animals was likely residual genomic material and not live and replicating virus. Therefore, SpFN was safe and effective at reducing disease and the amount of time that an animal was likely infectious following exposure to SARS-CoV-2.

Aside from determining the general efficacy of SpFN as a SARS-CoV-2 vaccine in CMs, this study also evaluated two different adjuvants: ALFQ and AlOH3. Although some differences in clinical disease characteristics existed between the two groups, most findings were minor or insignificant. However, there were some notable differences between these two groups, especially in terms of the immunological responses generated and disease in the lungs. The use of ALFQ was associated with higher neutralizing responses, including responses to the Alpha, Beta, and Gamma VoC and to the heterologous but related sarbecovirus SARS-CoV-1. Similar to the neutralizing antibody response, both ADCD and ADCP responses were stronger for the ALFQ adjuvant following initial vaccination; however, responses for both adjuvants were comparable after the boost vaccination. The use of ALFQ was also associated with a significantly higher pre- and post-challenge S1 subunit IgM response compared to AlOH3. The physiological relevance of a stronger IgM response was not elucidated in this study. However, the data suggest that the early antibody response following virus exposure may be stronger if ALFQ is used as the vaccine adjuvant.

The total IgA response was also significantly higher following virus challenge when ALFQ was used compared to AlOH3. IgA is a class of immunoglobulins most commonly responsible for protection of mucosal areas (including the respiratory tract and lungs) against infection [32]. For IgA, the difference in the response between Group 1 (ALFQ) and Group 2 (AlOH3) may have played a critical role in the control of disease at the predominant site of infection: the lungs. Group 2 radiographic findings in the lungs were of greater number compared to Group 1, and the lungs of Group 2 animals were slightly more affected pathologically compared to Group 1 animals. While findings in the lungs were less pronounced than those for the Group 3 controls, they demonstrated potential differences in efficacy based on adjuvant used.

Robust S-specific Th1 CD4+ T cell responses were elicited by SpFN adjuvanted with either ALFQ or AlOH3. Response magnitude did not significantly differ for most of the cytokine functions assessed, though mean values trended higher with ALFQ adjuvanting. IL-21 responses, a marker of T follicular helper CD4+ T cells, and greater T cell polyfunctionality were also notable with ALFQ.

SARS-CoV-2 variant introduction and spread in the human population has been a significant hindrance to the control of the ongoing pandemic; vaccines capable of protecting against current and emerging variants are a necessity. In this study, the ability of SpFN to generate a neutralizing antibody response against the Alpha, Beta, Gamma, Delta, and Omicron variants was assessed. Although there was significant reduction in neutralizing activity against VoCs compared to WA-1, with the exception of Omicron the Group 1 (SpFN + ALFQ) response to VoC was still relatively strong. This contrasts with the responses measured for Group 2 (SpFN + AlOH3) which were not only significantly lower than those against WA-1, but were also significantly lower for VoC compared to Group 1. Considering the robustness of these responses, it is predicted that vaccination with SpFN + ALFQ would be protective against infection with Alpha, Beta, Gamma, or Delta VoC.

Both SpFN + ALFQ and SpFN + AlOH3 were effective at reducing clinical disease following exposure of CMs to SARS-CoV-2. Overall, efficacy and strong immunogenicity have now been demonstrated in small animal models (mice and hamsters) and two nonhuman primate models (rhesus macaques and CM) of SARS-CoV-2. Use of the receptor-binding domain alone of the spike protein, adjuvanted with ALFQ, has also been highly protective in rhesus macaques [33]. Although SpFN + ALFQ performed better in these studies from an immunological perspective, SpFN + AlOH3 also promoted strong responses. The ability of SpFN to confer protection supports the need for follow-on studies to evaluate fractional vaccine doses, optimization of vaccination schedule, and additional comparative adjuvant dosing to assess the breadth and durability of protection against SARS-CoV-2 variants and related sarbecoviruses. Taken together, these data also support the evaluation of SpFN in clinical trials which are currently underway (Clinical trial number: NCT04784767 https://clinicaltrials.gov/ct2/show/NCT04784767).

## Materials and methods

### Animals

Animal research was conducted at the United States Army Medical Research Institute of Infectious Diseases (USAMRIID). Twenty-four adult *Macaca fascicularis* (cynomolgus macaques) of Asian origin were included on this study. Cynomolgus macaques were between 3-9 years old at time of challenge. Genders were mixed male and female, and all animals were SARS-CoV-2 naïve at the outset of the study. All animals passed a semi-annual physical examination and were certified as healthy by a veterinarian. Animals were acclimated in ABSL-3 animal rooms for 7 days prior to virus exposure and housed individually in 4.3 square foot cages. During the *in-life* portion of the study, animals were provided 2050 Monkey Chow (Harlan Teklad, Frederick, MD), fruits, and water *ad libitum* via an automatic watering system, and provided with enrichment regularly as recommended by the Guide for the Care and Use of Laboratory Animals.

### Ethics statement

These experiments and procedures were reviewed and approved by the United States Army Medical Research Institute for Infectious Diseases Institutional Animal Care and Use Committee (IACUC). All research was conducted in compliance with the USDA Animal Welfare Act (PHS Policy) and other federal statutes and regulations relating to animals and experiments involving animals, and adheres to the principles stated in the Guide for the Care and Use of Laboratory Animals, National Research Council, 2011. The facility is fully accredited by the Association for Assessment and Accreditation of Laboratory Animal Care, International. The animals were provided food and water *ad libitum* and checked at least once daily according to the protocol. All efforts were made to minimize painful procedures; the attending veterinarian was consulted regarding painful procedures, and animals were anesthetized prior to phlebotomy and virus infection. Animals were humanely euthanized at the end of study by intracardiac administration of a pentobarbital-based euthanasia solution under deep anesthesia in accordance with current American Veterinary Medical Association Guidelines on Euthanasia and institute standard operating procedures.

### Vaccine and vaccination

The SARS-CoV-2 SpFN nanoparticle vaccine contains the SARS-CoV-2 spike (S) glycoprotein (truncated at residue 158) linked to the *H. pylori* ferritin nanoparticle. Transfection, expression, and purification of SpFN was carried out as previously described [33-35]. In brief, an expression plasmid was transiently transfected into Expi293F cells using ExpiFectamine 293 transfection reagent and culture supernatants were harvested 4-5 days after transfection, filtered with a 0.22-µm filter. Protein was purified using GNA-lectin resin, followed by size-exclusion chromatography. Endotoxin levels were assessed (Endosafe® nexgen-PTS, Charles River Laboratories, Wilmington, MA) and 5 % v/v glycerol was added prior to filter-sterilization with a 0.22-µm filter, flash-freezing in liquid nitrogen, and storage at −80 °C. SpFN was thawed on the day of immunization at room temperature prior to adjuvant mixing. The SpFN immunogen was identical for Groups 1-2, with the only change being the adjuvant used. For Group 1, the vaccine was prepared at 50 µg in PBS containing ALFQ adjuvant [11.45 mM phospholipids (DMPC:DMPG=9:1), 55% cholesterol, 0.2 µg/ml 3D-PHAD (MPLA:PL=1:88), 0.1µg/ml QS-21]. For Group 2, the vaccine was prepared at 50 µg in PBS containing AlOH3 as the adjuvant. The concentration of the vaccine for administration was 50 µg/mL. On the day of vaccinations (Study Days -56 and -28), 1 mL of vaccine material (Groups 1-2) or PBS (Group 3) was administered to the deltoid. The vaccination site and day was the same for all animals, while alternating the deltoid side at the boost vaccination.

### Virus and virus exposure

A seed stock of SARS-CoV-2, isolate 2019-nCoV/USA-WA1/2020, designated as Lot R4719, was grown on ATCC Vero 76 cells using Stock Lot R4716 [Centers for Disease Control and Prevention (CDC) obtained]. The seed stock contains an average of 5.45×10^6^ pfu/mL of infectious virus as determined by plaque assay. R4719 was determined to have no detectable mycoplasma, endotoxin or adventitious agents based on the assays and techniques used. No known contaminants were detected when sequencing the stock. Identity was confirmed by real-time RT-PCR and sequencing. On the day of exposure (also called day of challenge), Study Day 1, animals were exposed to undiluted R4719 as follows: a total volume of 0.5 mL (0.25 mL per nare) was administered by the IN route, and 4.0 mL was administered by the IT route. The amount of virus in the challenge inoculum was determined by plaque assay.

### Animal observations and specimen collections

Animals were evaluated cage side for signs of illness. Other observations such as biscuit/fruit consumption, condition of stool, and urine output were also documented, if possible. Observations under anesthesia (physical examinations) occurred after cage side observations on Study Days -56, -42, -28, -14, -4, 1, 3, 5, 7, 9, 11, and 15. Pulse oximetry, radiography, blood collection, BAL collection, and collection of nasopharyngeal swab samples occurred during physical examinations. Nasopharyngeal swabs were collected into viral transport media (VTM; Hanks Balanced Salt Solution containing 2% heat-inactivated fetal bovine serum, 100 µg/mL gentamicin, and 0.5 µg/mL amphotericin B) and vortexed for 15-20 sec to create a homogenate. For preparation of sgRNA specimens, 500 µL of BAL fluid or swab homogenate was added to 500 µL ATL buffer. In addition, 100 uL of swab homogenate was added to 300 uL of TRIzol® LS (Thermo Fisher Scientific, Waltham, MA) for RNA isolation for RT-qPCR.

### Telemetry

Telemetry implants (M00; Data Sciences International, St. Paul, MN) were used to continuously monitor body temperature and activity in subject animals. Subjects were housed in individual cages in close proximity to radio frequency digital transceivers (TRX; Data Sciences International, St. Paul, MN) equipped with directional antennas pointed at the animal cages. These transceivers were connected via cat5e cables to a set of Communication Link Controllers (CLC, Data Sciences International, St. Paul, MN) to allow the digital multiplexing and the simultaneous collection of signals from all subjects. The signals were then routed over cat5e cable to data acquisition computers, which captured, reduced, and stored the digital data in data files (i.e., NSS files) using the Notocord-hem Evolution software platform (Version 4.3.0.77, Notocord Inc., Newark, New Jersey). Reduced data in the NSS files was extracted into Microsoft Excel workbooks using Notocord-derived formula add-ins, and the 30-minute (min) averages were calculated for each parameter for each subject. Telemetry data collected prior to challenge was used as baseline, and provided the average and standard deviation (SD) for each 30 min daily time of a 24-hour day.

### Necropsy, histology, and immunohistochemistry

Necropsies were conducted by a veterinary pathologist. The tissue samples were trimmed, routinely processed, and embedded in paraffin. Sections of the paraffin-embedded tissues 5 µm thick were cut for histology. For histology, slides were deparaffined, stained with hematoxylin and eosin (H&E), coverslipped, and labeled. Immunohistochemistry (IHC) was performed using the Dako Envision system (Dako Agilent Pathology Solutions, Carpinteria, CA, USA) After deparaffinization, peroxidase blocking, and antigen retrieval, replicate sections of lung, kidney, spleen and tracheobronchial lymph node were covered with a mouse monoclonal anti-SARS-CoV nucleocapsid protein (#40143-MM05, Sino Biological, Chesterbrook, PA, USA) at a dilution of 1:6000 and incubated at room temperature for forty five minutes. They were rinsed, and the peroxidase-labeled polymer (secondary antibody) was applied for thirty minutes. Slides were rinsed and a brown chromogenic substrate 3, 3’ Diaminobenzidine (DAB) solution (Dako Agilent Pathology Solutions) was applied for eight minutes. The substrate-chromogen solution was rinsed off the slides, and slides were counterstained with hematoxylin and rinsed. The sections were dehydrated, cleared with Xyless, and then cover slipped.

### Clinical pathology

For serum chemistries, whole blood was collected into Serum Clot Activator Greiner Vacuette tubes (Greiner Bio-One, Monroe, NC). Tubes were allowed to clot for at least 10 min and the serum separated in a centrifuge set at 1800 × g for 10 min at ambient temperature. The required volume of serum was removed for chemistry analysis using a General Chemistry 13 panel (Abaxis, Union City, CA) on a Piccolo Point-Of-Care Analyzer (Abaxis, Union City, CA). Serum was removed from the clot within 1 hour of centrifugation and was analyzed within 12 hours of collection.

For hematology, whole blood was collected into Greiner Vacuette blood tubes containing K3 EDTA as an anti-coagulant. Hematology was performed on VETSCAN® HM5 hematology analyzer (Abaxis, Union City, CA) within 4 hours of collection.

### sgRNA analysis

Viral RNA was extracted from 200 ul of nasopharyngeal swab and BAL material using the Qiagen EZ1 DSP Virus kit on the automated EZ1 XL Advance instrument (Qiagen, Valencia, CA). Real-time quantitative reverse transcription – polymerase chain reactions (RT-qPCR) were performed on the 7500 Dx Fast thermal cycler (Thermo Fisher Scientific, Life Technologies, Carlsbad, CA). Amplification and quantification of the sg RNA (targeting the E gene) was performed following the previously described method [33]. A synthetic RNA for subgenomic E was used as a calibrator. Results were reported in copies/ml.

### Plaque assay, RT-qPCR, Euroimmun SARS-CoV-2 S1 ELISA, and Magpix multiplex immunoassay

Plaque assay on challenge inoculum, RT-qPCR on TRIzol LS nasopharyngeal swab specimens, Euroimmun SARS-CoV-2 S1 ELISA, and Magpix multiplex immunoassay were all performed as described previously [20]. RT-qPCR on BAL material for amplification and quantitation of viral RNA (targeting the E gene) was performed as described above for sgRNA analysis.

### Biolayer interferometry, PRNT, and pseudovirion assays

PRNT was performed as described previously [20]. The viruses used in the PRNT were SARS-CoV-2, isolate 2019-nCoV/USA-WA1/2020, designated as Lot R4719 described above. The other three VoC used in the PRNT were Beta (Catalog Number NR-54009 contributed by Alex Sigal and Tulio de Oliveira), Gamma (Catalog Number NR-54984), and Delta (Catalog Number NR-55674 contributed by Dr. Andrew Pekosz) obtained through BEI Resources, NIAID, NIH.

RBD molecules for WA-1 and VoC were produced, and the biolayer interferometry assays were performed as previously described [33-35]. In brief, ForteBio HIS1K biosensors were hydrated in PBS prior to use. All assay steps were performed at 30°C with agitation set at 1,000 rpm in the Octet RED96 instrument (FortéBio). His-tagged RBD molecules (30 μg/ml diluted in PBS) were allowed to load on the probes for 120 seconds. After briefly dipping in assay buffer (15 seconds in PBS), the biosensors were dipped in NHP sera (50-fold dilution) for 180 seconds. Binding response (nm) was reported for the 180s timepoint.

Pseudovirion assays against SARS-CoV-1 and SARS-CoV-2 were performed as follows. The S expression plasmid sequence for SARS-CoV-2 was codon optimized and modified to remove an 18 amino acid endoplasmic reticulum retention signal in the cytoplasmic tail to improve S incorporation into pseudovirions (PSV) and thereby enhance infectivity. SARS-CoV-2 pseudovirions (PSV) were produced by co-transfection of HEK293T/17 cells with a SARS-CoV-2 S plasmid (pcDNA3.4), derived from the Wuhan-Hu-1 genome sequence (GenBank accession number: MN908947.3) and an HIV-1 (pNL4-3.Luc.R-E-, NIH HIV Reagent Program, Catalog number 3418). S expression plasmids for SARS-CoV-2 VOC were similarly codon optimized and modified and included the following mutations: B.1.1.7 (69-70del, Y144del, N501Y, A570D, D614G, P681H, T718I, S982A, D1118H), B.1.351 (L18F, D80A, D215G, 241-243del, K417N, E484K, N501Y, D614G, A701V, E1195Q). Infectivity and neutralization titers were determined using ACE2-expressing HEK293 target cells (Integral Molecular, Philadelphia, PA) in a semi-automated assay format using robotic liquid handling (Biomek NXp Beckman Coulter, Brea, CA). Virions pseudotyped with the vesicular stomatitis virus (VSV) G protein were used as a non-specific control. Test sera were diluted 1:40 in growth medium and serially diluted; then 25 μl/well was added, in triplicate, to a white 96-well plate. An equal volume of diluted SARS-CoV-2 PSV was added to each well and plates were incubated for 1 hour at 37°C. Target cells were added to each well (40,000 cells/ well) and plates were incubated for an additional 48 hours. Relative light units (RLU) were measured with the EnVision Multimode Plate Reader (Perkin Elmer, Waltham, MA) using the Bright-Glo Luciferase Assay System (Promega, Madison, WI). Neutralization dose–response curves were fitted by nonlinear regression using the LabKey Server, as previously described [36]. Final titers are reported as the reciprocal of the dilution of serum necessary to achieve 50% (ID50, 50% inhibitory dose) and 90% neutralization (ID90, 90% inhibitory dose). Assay equivalency was established by participation in the SARS-CoV-2 Neutralizing Assay Concordance Survey (SNACS) run by the Virology Quality Assurance Program and External Quality Assurance Program Oversite Laboratory (EQAPOL) at the Duke Human Vaccine Institute, sponsored through programs supported by the National Institute of Allergy and Infectious Diseases, Division of AIDS.

### Serum binding and ACE-2 inhibitory antibody assessment

SARS-CoV-2-specific binding IgG antibody responses were measured using MULTI-SPOT^®^ 96-well plates, V-PLEX SARS-CoV-2 Panel 7 Kit (Meso Scale Discovery (MSD), Rockville, MD). Multiplex wells were coated with three SARS-CoV-2 antigens, S, RBD, and Nucleocapsid (N) from WA-1, and S and RBD from Alpha (B.1.1.7), Beta (B.1.351) and Gamma (P.1) at a concentration of 200-400 ng/ml. Bovine serum albumin (BSA) served as a negative control (background signal). 10-plex MULTISPOT plates were blocked with MSD Blocker A buffer for 1 hour at room temperature (RT) while shaking at 700 rpm. Plates were washed with buffer before the addition of reference standard, calibrator controls, and samples. Serum/plasma samples were diluted at 1:1,000 - 1:200,000 in diluent 100 buffer, then added to duplicate wells. Plates were incubated for 2 hours at RT while shaking at 700 rpm, then washed. MSD SULFO-TAG^™^ conjugated anti-IgG antibody was added to each well. Plates were incubated for 1 hour at RT with shaking at 700 rpm and washed, then MSD GOLD^™^ Read buffer was added to each well. Plates were read by the MESO SECTOR S600 Reader. IgG concentration was calculated using DISCOVERY WORKBENCH^®^ MSD Software and converted to Binding Antibody Units (BAU/mL) using the WHO/NIBSC standard.

The ability of SARS-CoV-2 Spike-specific binding antibodies to inhibit S or RBD binding to the ACE-2 receptor was also measured using the same V-PLEX SARS-CoV-2 Panel 7 Kit MULTI-SPOT^®^ 96-well plates (MSD, Rockville, MD) but with ACE2 protein conjugated with MSD SULFO-TAG^™^ in a competition format. Antigen-coated plates were blocked and washed as described above. Assay calibrator and samples were diluted at 1:25 - 1:1,000 in MSD Diluent 100 buffer, then added to the wells. Plates were incubated for 1 hour at RT while shaking at 700 rpm. ACE2 protein conjugated with MSD SULFO-TAG^™^ was added, and plates were incubated for 1 hour at RT while shaking at 700rpm. Plates were washed and read as described above. AU/mL concentration of inhibitory antibodies was calculated with DISCOVERY WORKBENCH^®^ MSD Software.

### ADCP and ADCD assays

ADCP was measured as previously described [37]. Briefly, biotinylated SARS-CoV-2 Spike trimer (Hexapro) was incubated with red streptavidin-fluorescent beads (Molecular Probes, Eugene, OR) for 2h at 37°C. 10 μl of a 100-fold dilution of beads–protein was incubated 2h at 37°C with 100μl 900-fold diluted plasma samples before addition of THP-1 cells (25,000 cells per well; Millipore Sigma, Burlington, MA). After 19h incubation at 37°C, the cells were fixed with 2% formaldehyde solution (Tousimis, Rockville, MD) and fluorescence was evaluated on a LSRII (BD Bioscience, San Jose, CA). The phagocytic score was calculated by multiplying the percentage of bead-positive cells by the geometric mean fluorescence intensity (MFI) of the bead-positive cells and dividing by 10^4^.

An ADCD assay was adapted from [38]. Briefly, SARS-CoV-2 Spike-expressing 293 FreeStyle (293F) cells were generated by transfection with linearized plasmid (pcDNA3.1) encoding codon-optimized full-length SARS-CoV-2 Spike protein matching the amino acid sequence of the IL-CDC-IL1/2020 isolate (GenBank ACC# MN988713). Stable transfectants were single-cell sorted and selected to obtain a high-level Spike surface expressing clone (293F-Spike-S2-WT). 293F-Spike-S2-WT cells were incubated with 10-fold diluted heat-inactivated (56°C for 30 min) plasma samples for 30 min at 37°C. Cells were washed twice and resuspended in R10 media. Lyophilized guinea pig complement (CL4051, Cedarlane, Burlington, Canada) was reconstituted per the manufacturer’s instructions in 1 mL cold water and centrifuged for 5 min at 4°C to remove aggregates. Cells were washed with PBS and resuspended in 200 µl of guinea pig complement, which was prepared at a 1:50 dilution in Gelatin Veronal Buffer with Ca2+ and Mg2+ (IBB-300x, Boston BioProducts, Ashland, MA). After incubation at 37°C for 20 min, cells were washed in PBS 15mM EDTA (ThermoFisher Scientific, Waltham, MA) and stained with an anti-guinea pig complement C3 FITC (polyclonal, ThermoFisher Scientific, Waltham, MA). Cells were fixed with 4% formaldehyde solution and fluorescence was evaluated on a LSRII (BD Bioscience, San Jose, CA).

### Antigen-specific T cell responses

Cryopreserved peripheral blood mononuclear cells were thawed and rested for 6 h in R10 supplemented with 50 U/mL Benzonase Nuclease (Sigma-Aldrich, St. Louis, MO) at 37^°^C followed by stimulation with peptide pools for 12 h. Stimulations consisted of two pools of peptides spanning the S protein of SARS-CoV-2 or SARS-CoV-1 (1 µg/mL, JPT, PM-WCPV-S and PM-CVHSA-S respectively) in the presence of Brefeldin A (0.65 µL/mL, GolgiPlug^™^, BD Cytofix/Cytoperm Kit, Cat. 555028), co-stimulatory antibodies anti-CD28 (BD Biosciences Cat. 555725 1 µg/mL) and anti-CD49d (BD Biosciences Cat. 555501; 1ug/mL) and CD107a (H4A3, BD Biosciences Cat. 561348, Lot 9143920 and 253441). Following stimulation, cells were stained serially with LIVE/DEAD Fixable Blue Dead Cell Stain (ThermoFisher #L23105) and a cocktail of fluorescent-labeled antibodies (BD Biosciences unless otherwise indicated) to cell surface markers CD4-PE-Cy5.5 (S3.5, ThermoFisher #MHCD0418, Lot 2118390 and 2247858), CD8-BV570 (RPA-T8, BioLegend #301038, Lot B281322), CD45RA BUV395 (5H9, #552888, Lot 154382 and 259854), CD28 BUV737 (CD28.2, #612815, Lot 0113886), CCR7-BV650 (GO43H7, # 353234, Lot B297645 and B316676) and HLA-DR-BV480 (G46-6, # 566113, Lot 0055314). Intracellular cytokine staining was performed following fixation and permeabilization (BD Cytofix/Cytoperm, BD Biosciences) with CD3-Cy7APC (SP34-2, #557757, Lot 6140803 and 121752), CD154-Cy7PE (24-31, BioLegend # 310842, Lot B264810 and B313191), IFN*γ*-AF700 (B27, # 506516, Lot B187646 and B290145), TNF*α*-FITC (MAb11, # 554512, Lot 15360), IL-2-BV750 (MQ1-17H12, BioLegend #566361, Lot 0042313), IL-4 BB700 (MP4-25D2, Lot 0133487 and 0308726), MIP-1b (D21-1351, # 550078, Lot 9298609), CD69-ECD (TP1.55.3, Beckman Coulter # 6607110, Lot 7620070 and 7620076), IL-21-AF647 (3A3-N2.1, # 560493, Lot 9199272 and 225901), IL-13-BV421 (JES10-5A2, # 563580, Lot 9322765, 210147 and 169570) and IL-17a-BV605 (BL168, Biolegend #512326, B289357). Sample staining was measured on a FACSymphony(tm) A5 SORP (Becton Dickenson) and data was analyzed using FlowJo v.9.9 software (Tree Star, Inc.). CD4+ and CD8+ T cell subsets were pre-gated on memory markers prior to assessing cytokine expression as follows: single-positive or double-negative for CD45RA and CD28. Boolean combinations of cells expressing one or more cytokines were used to assess the total S-specific response of memory CD4+ or CD8+ T cells. Responses from the two-peptide pools spanning SARS-CoV-2 S or SARS-CoV-1 S were summed. Values were background corrected by subtraction of the unstimulated (DMSO) condition, and responses three times higher than the group average prior to immunization (study Day -56) were considered positive. Display of multicomponent distributions were performed with SPICE v6.0 (NIH, Bethesda, MD).

The COMPASS method for determination of T cell functionality and polyfunctionality was performed as described previously [30]. Functionality score represents the proportion of antigen-specific subsets detected among all possible combinations of subsets. The polyfunctionality score is similar to the functionality score but weighs the different subsets by their degree of functionality, statistically favoring subsets with higher degrees of functionality. Representative FACS staining and gating strategy for T cell intracellular cytokine staining can be found in S1 Fig.

## Acknowledgements

The authors would like to acknowledge the members of the United States Army Medical Research Institute of Infectious Diseases (USAMRIID) Veterinary Medicine Division and the Core Laboratory Sciences Division for their contributions during protocol conception and study conduct. Specifically, the authors would like to thank the following individuals from those divisions for their efforts: MAJ Philip Bowling, Joshua Shamblin, Joshua Moore, Jimmy Fiallos, Willie Sifford, Eugene Blue, Stephen Stevens, SGT Robin Cornelius, Emily Cornelius, David Dyer, Ondraya Frick, Kerry Berrier, Christopher Jensen, David Nyakiti, Jeanean Ghering, Nazira (Ashley) Alli, Heather Esham, Holly Bloomfield, Stephanie Bellanca, Cristal Johnson, Joshua Johnson, Mary Leyva, Joshua McClain, and Brandon Tapia. The authors also thank the following Walter Reed Army Institute of Research (WRAIR) members for their superb technical support: Ming Dong, Viviana Cobos Jimenez, Dante Coleman, Hannah Grove, Heather Hernandez, and MAJ Kathryn McGuckin Wuertz. Finally, the authors would like to thank the program manager, Kristan O’Brien, for her assistance with programmatic requirements including funding execution, timeline management, and deliverable tracking.

## Disclaimer

### USAMRIID Disclaimer

Opinions, interpretations, conclusions, and recommendations are those of the author and are not necessarily endorsed by the U.S. Army. Funding for this effort was provided by the Military Infectious Diseases Research Program under project number 341155776.

### WRAIR Disclaimer

The opinions or assertions contained herein are the private views of the authors, and are not to be construed as official, or as reflecting true views of the Department of the Army or the Department of Defense or HJF. This work was supported by a cooperative agreement (W81XWH-18-2-0040) between the Henry M. Jackson Foundation for the Advancement of Military Medicine, Inc. and the U.S. Department of Defense (DOD), and was supported in part by the US Army Medical Research and Development Command under contract number No. W81-XWH-18-C-0337.

## Supporting information

**S1 Fig. Representative FACS staining and gating strategy for T cell ICS**. Flow cytometry gating applied to macaque PBMC T cell ICS data. FACS plots depict staining following stimulation with DMSO (A) or SARS-CoV-2 spike peptide pool #1 (B) for a representative animal at study week 6 following two immunizations with 50 μg SpFN adjuvanted with ALFQ. Sequential gates were applied from left to right (top row) to identify CD4+ and CD8+ T cells. Cytokine production was measured in total memory CD4+ (middle) and CD8+ (bottom) T cells by excluding the naïve CD28+ CD45RA+ population. Cytokine-positive cells were identified by co-expression of CD69.

**S2 Fig. SARS-CoV-2 and SARS-CoV-1 S-specific CD8 and CD4 T cell responses elicited by adjuvanted SpFN vaccination**. T cell responses were assessed by SARS-CoV-2 or SARS-CoV-1 spike peptide pool stimulation and intracellular cytokine staining of PBMC collected at study day -56 (pre-immune), study day -42 (2 weeks post-prime) and study day -14 (2 weeks post-boost). (A) SARS-CoV-2 S-specific memory CD8+ T cells expressing one or more Th1 cytokines (IFN-*γ*, TNF-*α*, and IL-2) is shown. (B) SARS-CoV-1 S-specific memory CD4+ T cells expressing IL-21 is shown. The fraction of animals within each group with a positive response following each vaccination is indicated. Significant differences between groups was assessed using a Kruskal-Wallis test followed by a Dunn’s post-test (*<0.05, **<0.01).

**S1 Table. Summary of clinical disease findings**

**S2 Table. Summary of clinical pathology findings – VP**

**S3 Table. Summary of clinical pathology findings – CP**

**S1 Appendix. Summary of histologic findings (excluding the lungs and nasal turbinates)**

**S2 Appendix. Summary of major histopathologic findings in the respiratory tract**

