## Supplemental Files for "A SARS-CoV-2 Spike Ferritin Nanoparticle Vaccine is Protective and Promotes a Strong Immunological Response in the Cynomolgus Macaque Coronavirus Disease 2019 (COVID-19) Model"

### Supplementary Materials for

Title: Efficacy Evaluation of Novel SARS-CoV-2 Vaccines in a Coronavirus Disease 2019 (COVID-19) Nonhuman Primate Model

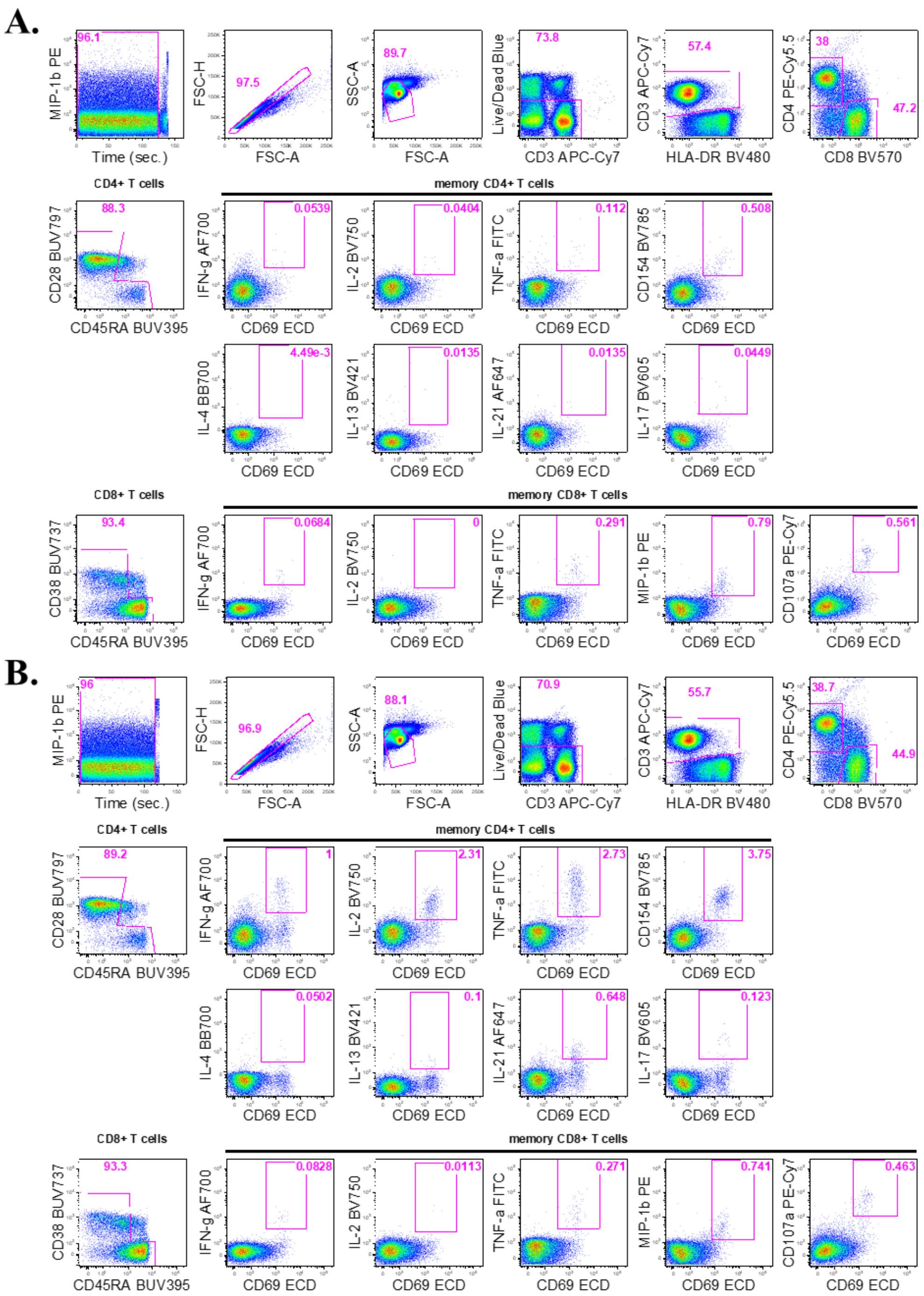


**S1 Fig. Representative FACS staining and gating strategy for T cell ICS.** Flow cytometry gating applied to macaque PBMC T cell ICS data. FACS plots depict staining following stimulation with DMSO (A) or SARS-CoV-2 spike peptide pool #1 (B) for a representative animal at study week 6 following two immunizations with 50 µg SpFN adjuvanted with ALFQ. Sequential gates were applied from left to right (top row) to identify CD4+ and CD8+ T cells. Cytokine production was measured in total memory CD4+ (middle) and CD8+ (bottom) T cells by excluding the naïve CD28+ CD45RA+ population. Cytokine-positive cells were identified by co-expression of CD69.

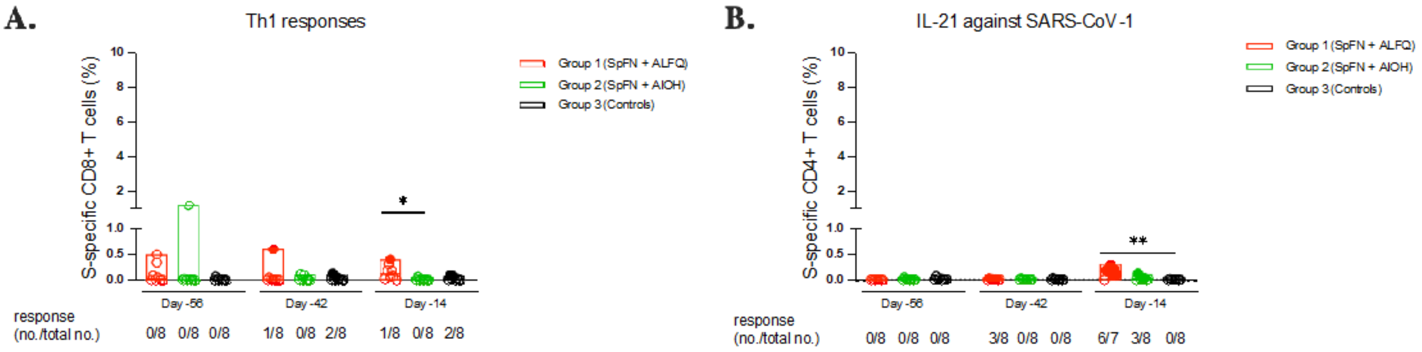


**S2 Fig**. **SARS-CoV-2 and SARS-CoV-1 S-specific CD8 and CD4 T cell responses elicited by adjuvanted SpFN vaccination.** T cell responses were assessed by SARS-CoV-2 or SARS-CoV-1 spike peptide pool stimulation and intracellular cytokine staining of PBMC collected at study day -56 (pre-immune), study day -42 (2 weeks post-prime) and study day -14 (2 weeks post-boost). (A) SARS-CoV-2 S-specific memory CD8+ T cells expressing one or more Th1 cytokines (IFN-γ, TNF-α, and IL-2) is shown. (B) SARS-CoV-1 S-specific memory CD4+ T cells expressing IL-21 is shown. The fraction of animals within each group with a positive response following each vaccination is indicated. Significant differences between groups was assessed using a Kruskal-Wallis test followed by a Dunn's post-test (*<0.05, **<0.01).

#### S1 Table. Summary of clinical disease findings

| CM # | 2 | 5 | 7 | 8 | 9 | 14 | 21 | 24 | 1 | 3 | 4 | 6 | 10 | 16 | 18 | 23 | 11 | 12 | 13 | 15 | 17 | 19 | 20 | 22 |
|---|---|---|---|---|---|---|---|---|---|---|---|---|---|---|---|---|---|---|---|---|---|---|---|---|
| Group | 1 | 1 | 1 | 1 | 1 | 1 | 1 | 1 | 2 | 2 | 2 | 2 | 2 | 2 | 2 | 2 | 3 | 3 | 3 | 3 | 3 | 3 | 3 | 3 |
| Euthanasia Day | 15 | 15 | 15 | 15 | 9 | 9 | 9 | 9 | 15 | 9 | 15 | 15 | 15 | 9 | 9 | 9 | 9 | 9 | 9 | 15 | 9 | 15 | 15 | 15 |
| Hyperpyrexia[1] | | | | | | | | | | | | | | | | | | | X | | X | | X | |
| Fever[2] | | | | | X | X | | | | | | | | | | | | X | X | X | X | X | X | |
| Significantly Elevated Body Temp[3] | X | | | | X | X | | X | | X | | | | | | | X | X | X | X | X | X | X | X |
| Diurnal Rhythm Disruption[4] | X | | X | | | X | | | | X | | X | | | | | | | | | | | | |
| Mild Hypoxia[5] | | | | | | X | | X | | | | X | | | | | | | | | | | X | X |
| Tachycardia | | X | X | X | X | X | X | X | | | X | X | X | X | X | X | | X | X | X | X | X | X | X |
| Cough | | | | X | X | | | | | X | | | | | | | | X | | | X | | | |
| Increased Lung Opacity (# Affected Lobes) | X (1) | | X (1) | X (1)^ | | | X (1)^ | | | X (4) | X (1) | | | | X (2) | | | | X (2) | X (2+) | X (4) | | | |
| Lung Infiltrates (# Affected Lobes) | | | | | | | | | X (1) | X (2) | | | | | | | X (1) | | | X (1) | X (4) | X (2) | | |
| Abdominal Component to Breathing | | X | | | | X | | | | | | X | | | | | X | | | | X | | | |
| Piloerection | | | | | | | | X | X | X | | X | | X | X | | X | X | | X | X | | | |
| Decreased Skin Turgor | | | | | | | | | X | | | | X | | | | | | | X | | X | | X |
| Lymphadenopathy | | | | X | | | | | | | | | | | | | | | | X | | X | | |
| Stool Not Fully Formed/Liquid | | | X | X | X | | | | | X | | X | X | | | | | | | | | | | |
| Anorexia[6] | | X | | X | | | | | X | | X | X | | | | X | X | X | | X | X | | | |
| Reduced Consumption[7] | | | | X | X | | | | X | X | X | X | | X | | X | X | X | | X | X | | | |

*Mild hypoxia is defined as 90-94% SpO2

[1]Defined as a temperature greater than or equal to 3.0°C above baseline by telemetry, with baseline being the mean of data from Study Days -7 through -1 for a particular animal

[2]Defined as a temperature greater than or equal to 1.5°C above baseline for a duration of greater than 2 hours by telemetry

[3]Defined as a temperature greater than 3 standard deviations above baseline for a duration of greater than 2 hours by telemetry.

[4]Defined as a significant activity increase during the dark cycle, or a significant activity decrease during the light cycle, compared to baseline as measured by telemetry

[5]Defined as 90-94% $SpO_2$

[6]Defined as an absence of biscuit and enrichment consumption for one or more days OR an absence of biscuit consumption for 3 or more consecutive days

[7]Defined as either no evidence or notably reduced amounts of biscuit and/or fruit consumption for greater than or equal to 2 consecutive days

^Potential positional or rotational artifact

#### S2 Table. Summary of clinical pathology findings – VP

| CM # | 2 | 5 | 7 | 8 | 9 | 14 | 21 | 24 | 1 | 3 | 4 | 6 | 10 | 16 | 18 | 23 | 11 | 12 | 13 | 15 | 17 | 19 | 20 | 22 |
|---|---|---|---|---|---|---|---|---|---|---|---|---|---|---|---|---|---|---|---|---|---|---|---|---|
| Group | 1 | 1 | 1 | 1 | 1 | 1 | 1 | 1 | 2 | 2 | 2 | 2 | 2 | 2 | 2 | 2 | 3 | 3 | 3 | 3 | 3 | 3 | 3 | 3 |
| ↑ WBC | | | | | | | | | | | X | | | | | | | | | | | | | |
| ↑ NEUT | | | | | | | | | | | X | | | X | | | | | | | | | | |
| ↑ EOS | | | | | | | | | | | X | | | | | | | | | | | | | |
| ↑ BAS | | | | | | | | | | | X | | | | | | | | | | | | | |
| ↑ LYM | | X | | | | | | X | | | | | | | | | | | X | | | | | X |
| ↑ MON | | X | | | | | | | | | | | | | | | | | X | | | | X | |
| ↑ ALT | | X | | | | | | | | X | | | | | | X | | X | | | | | | |
| ↓ ALB | | | | | | X | | | | | | | | | | | | | | | | | | |
| ↑ ALP | | | | | | | | | | | X | | | | | | | | | | | | | |
| ↑ AST | | X | | | | | | X | | | | | | | | | | | | | | | X | |
| ↑ AMY | | | | X | | | | | | | | | | | | | | | | | | | | |
| ↑ BUN | | X | | | | | | | | | | | | | | | X | | | | | | | |
| ↑ CRE | | | | | | | X | | | | | | | | | | | | | | | | | |
| ↑ TBIL | | | | | | | | | | | | | | X | | | | | | | | | | |
| ↑ CK | | X | X | | | X | | X | | X | X | X | X | | X | X | | | X | X | | X | X | |
| ↑ CRP | | | | | | X | | | | | | | | | | | | | | | | | | |

X = % change from baseline (Study Day -56) of ≥30%

30-49% Δ

50-69% Δ

70-99% Δ

100-499% Δ

>500%

##### S3 Table. Summary of clinical pathology findings – CP

| CM # | 2 | 5 | 7 | 8 | 9 | 14 | 21 | 24 | 1 | 3 | 4 | 6 | 10 | 16 | 18 | 23 | 11 | 12 | 13 | 15 | 17 | 19 | 20 | 22 |
|---|---|---|---|---|---|---|---|---|---|---|---|---|---|---|---|---|---|---|---|---|---|---|---|---|
| Group | 1 | 1 | 1 | 1 | 1 | 1 | 1 | 1 | 2 | 2 | 2 | 2 | 2 | 2 | 2 | 2 | 3 | 3 | 3 | 3 | 3 | 3 | 3 | 3 |
| ↑ WBC | | | | | X | | X | | X | X | X | X | | X | | | | X | | X | | X | X | X |
| ↑ NEUT | X | | | | X | | X | X | X | X | X | X | | X | X | | | | X | X | | X | X | X |
| ↑ EOS | X | | X | X | X | | X | | X | | X | X | | X | X | | | | | X | X | X | X | X |
| ↑ BAS | | | | | X | | | | X | | X | X | | X | | | | | | X | | X | X | X |
| ↑ LYM | | X | X | X | | X | X | | X | | X | X | X | | | | | X | | X | | X | | X |
| ↑ MON | | X | X | X | X | X | | X | X | X | X | | X | | X | | X | X | | X | X | X | X | X |
| ↓ PLT | | | X | X | | | | | | | X | | | | | X | | | | | | | | X |
| ↑ ALT | | | X | X | | | | | | | | | | X | | X | | X | X | X | | | | |
| ↑ ALP | | | | | | | | X | | | | | | | | | X | X | X | X | | X | | X |
| ↑ AST | X | | X | | | | | X | X | | X | X | X | X | | X | X | X | X | X | X | X | | |
| ↑ GGT | | | | | | | | | | | | | | | | | | | | | | | | X |
| ↑ GLU | | | X | | | | | X | | | | | | | | | | | | | | | | |
| ↑/↓ AMY | | | | | | | | X | | | | | | | | | X | | | | | | X | |
| ↑ BUN | X | | | X | | | | | | | | | | | | | | | | | | | | |
| ↑ CRE | | X | X | | | X | | | | | | | | | | | | | | X | | | | |
| ↑ TBIL | | | | | | | | X | | | X | | X | | | | X | | | X | | | X | |
| ↑ CK | X | X | X | X | X | X | X | X | X | X | X | X | X | X | X | X | | X | X | X | X | X | X | |
| ↑ CRP | | X | X | X | X | X | | X | | | | X | | | | | X | X | X | X | X | X | X | |

X = % change from baseline (average of Study Days -4 and 1 per animal) of ≥30%

30-49% Δ

50-69% Δ

70-99% Δ

100-499% Δ

500-999% Δ

>1000% Δ

#### S1 Appendix. Summary of histologic findings (excluding the lungs and nasal turbinates)

| Animal Number | Group | Tracheobronchial Lymph Node | Axillary Lymph Node | Inguinal Lymph Node | Liver | GI Tract | Other |
|---|---|---|---|---|---|---|---|
| CM 2 | 1 | Moderate lymphoid hyperplasia | Mild lymphoid hyperplasia | Medullary fibrosis | Hepatocyte atrophy, capsular fibrosis | Lymphoid hyperplasia of the ileocecolic lymph node | Testicular atrophy, urinary bladder and prostate gland infiltrate, interstitial nephritis |
| CM 5 | 1 | Moderate lymphoid hyperplasia | Minimal lymphoid hyperplasia | Draining hemorrhage | | Stomach inflammation | Urinary bladder infiltrate, ovarian cyst, skeletal muscle degeneration, necrosis and regeneration with inflammation |
| CM 7 | 1 | | Medullary fibrosis | Medullary fibrosis | | Stomach inflammation | Splenic lymphoid hyperplasia, prostate gland infiltrate |
| CM 8 | 1 | Mild lymphoid hyperplasia | | | | Stomach inflammation | Adrenal gland mineralization |
| CM 9 | 1 | Minimal lymphoid hyperplasia | Medullary fibrosis | | | | Heart, trachea and prostate infiltrate |
| CM 14 | 1 | Mild lymphoid hyperplasia | | | | | Interstitial nephritis, ovarian corpus amalacea |
| CM 21 | 1 | Moderate lymphoid hyperpasia, medullary infiltrate | | Medullary fibrosis | | Stomach inflammation | Tracheal infiltrate, mesenteric lymphoid hyperplasia |
| CM 24 | 1 | | Granulomatous inflammation | Granulomatous inflammation | | Cecal parasite | |
| CM 1 | 2 | Marked lymphoid hyperplasia | | | | Stomach inflammation, lymphoid hyperplasia ileocecolic LN | Splenic lymphoid hyperplasia, interstitial nephritis |
| CM 3 | 2 | Mild lymphoid hyperplasia | | | Lymphocytic infiltrate | | Splenic lymphoid hyperplasia |
| CM 4 | 2 | Mild lymphoid hyperplasia | Medullary fibrosis | | Lymphocytic infiltrate | Stomach inflammation, GALT lymphoid hyperplasia | Splenic lymphoid hyperplasia, tracheal infiltrate |

| Animal Number | Group | Tracheobronchial Lymph Node | Axillary Lymph Node | Inguinal Lymph Node | Liver | GI Tract | Other |
|---|---|---|---|---|---|---|---|
| CM 6 | 2 | | Mild lymphoid hyperplasia | | Lymphocytic infiltrate | Stomach inflammation, lymphoid hyperplasia esophagus, duodenal infiltrate | Ovarian corpora amylacea |
| CM 10 | 2 | | Medullary fibrosis | Medullary fibrosis, draining hemorrhage | | Peyer's patch lymphoid hyperplasia | Infiltrate in heart, urinary bladder and prostate gland; interstitial nephritis |
| CM 16 | 2 | | Reticuloendo-thelial cell hyperplasia | Medullary fibrosis, draining hemorrhage | | | Lymphoid hyperplasia in kidney, prostate gland infiltrate |
| CM 18 | 2 | Lymphoid hyperplasia, medullary infiltrate | Lymphoid hyperplasia | Medullary fibrosis | | Ileal and cecal parasite | Infiltrate in heart, prostate gland and sciatic nerve |
| CM 23 | 2 | | | | | Stomach inflammation | Ovarian mineralization |
| CM 11 | 3 | | Minimal lymphoid hyperplasia | | | | Skeletal muscle regeneration |
| CM 12 | 3 | | | Draining hemorrhage | | Stomach inflammation | Skeletal muscle degeneration & necrosis with inflamamtion |
| CM 13 | 3 | Mild lymphoid hyperplasia | | | | | Thyroid gland infiltrate |
| CM 15 | 3 | Mild lymphoid hyperplasia | | | Neutrophilic and eosinophilic infiltrate | Esophageal and cecal infiltrate, lymphoid hyperplasia ileocecolic LN | Heart degeneration with inflammation, urinary bladder and kidney infiltrate, |
| CM 17 | 3 | Mild lymphoid hyperplasia | | | Mononuclear infiltrate | Stomach inflammation, cecal parasite | Prostate gland infiltrate |
| CM 19 | 3 | Mild lymphoid hyperplasia | | | | | Interstitial nephritis |
| CM 20 | 3 | Minimal lymphoid hyperplasia | | | | | Infiltrate in urinary bladder and corpus striatum |
| CM 22 | 3 | Minimal lymphoid hyperplasia | | Medullary fibrosis | | Infiltrate in esophagus, cecal parasite | Heart infiltrate, interstitial nephritis |

#### S2 Appendix. Summary of Major Histopathologic Findings in the Respiratory Tract

| Animal Number | Group | Alveolar inflammation | Perivascular inflammation | Peribronchiolar inflammation | Pleural inflammation and/or fibrosis | Nasal turbinate inflammation |
|---|---|---|---|---|---|---|
| CM 2 | 1 | None | Mild to moderate; 2 of 8 sections affected | None | None | Minimal; 2 of 6 sections affected |
| CM 5 | 1 | None | None | None | None | None |
| CM 7 | 1 | None | None | None | None | None |
| CM 8 | 1 | None | Minimal/1 of 8 sections affected | None | None | None |
| CM 9 | 1 | Minimal; 1 of 8 sections affected | Minimal to mild; 6 of 8 sections affected | Mild; 2 of 8 sections affected | Mild; 2 of 8 sections affected | Minimal to mild; 2 of 6 sections affected |
| CM 14 | 1 | Minimal/1 of 8 sections affected | Minimal to moderate; 3 of 8 sections affected | Mild/2 of 8 sections affected | None | None |
| CM 21 | 1 | Minimal/2 of 8 sections affected | Minimal/3 of 8 sections affected | None | None | Minimal/1 of 6 sections affected |
| CM 24 | 1 | None | None | None | None | Minimal; 1 of 6 sections affected |
| CM 1 | 2 | None | Minimal/1 of 8 sections affected | None | Minimal/1 of 8 sections affected | Minimal - mild; 3 of 6 sections affected |
| CM 3 | 2 | None | Minimal/2 of 8 sections affected | None | Minimal/1 of 8 sections affected | None |
| CM 4 | 2 | Minimal; 1 of 8 sections affected | None | None | Mild; 1 of 8 sections affected | None |
| CM 6 | 2 | Mild; 1 of 8 sections affected | Minimal - mild; 2 of 8 sections affected | None | Mild/1 of 8 sections affected | Mild - moderate; 3 of 6 sections affected |

| Animal Number | Group | Alveolar inflammation | Perivascular inflammation | Peribronchiolar inflammation | Pleural inflammation and/or fibrosis | Nasal turbinate inflammation |
|---|---|---|---|---|---|---|
| CM 10 | 2 | None | None | None | None | None |
| CM 16 | 2 | Minimal to mild; 2 of 8 sections affected | Minimal to moderate; 4 of 8 sections affected | Mild to moderate; 2 of 8 sections affected | Mild; 2 of 8 sections affected | None |
| CM 18 | 2 | None | Minimal to mild; 2 of 8 sections affected | Mild; 1 of 8 sections affected | None | None |
| CM 23 | 2 | None | None | None | None | Minimal/1 of 6 sections affected |
| CM 11 | 3 | Minimal - moderate; 7 of 8 sections affected | Mild - moderate; 2 of 8 sections affected | Mild - moderate; 3 of 8 sections affected | None | None |
| CM 12 | 3 | Minimal to mild; 4 of 8 sections affected | Minimal to mild; 4 of 8 sections affected | None | None | None |
| CM 13 | 3 | Moderate/1 of 8 sections affected | Minimal - moderate; 5 of 8 sections affected | Mild/2 of 8 sections affected | None | Minimal/1 of 6 sections affected |
| CM 15 | 3 | Mild; 1 of 8 sections affected | Moderate; 1 of 8 sections affected | None | None` | Mild-moderate; 5 of 6 sections affected |
| CM 17 | 3 | Mild; 2 of 8 sections affected; fibrin present | Mild; 2 of 8 sections affected | None | Mild/1 of 8 sections affected | None |
| CM 19 | 3 | Minimal to mild; 2 of 8 sections affected | Minimal - mild; 4 of 8 sections affected | None | None | Minimal; 2 of 6 sections affected |
| CM 20 | 3 | Mild; 1 of 8 sections affected | Mimial to mild; 4 of 8 sections affected | None | None | Minimal - 1 of 6 sections affected |
| CM 22 | 3 | None | Minimal to mild; 2 of 8 sections affected | None | None | None |
